# Targeting AASS improves neurotoxicity and mitochondrial function in astrocyte models for pyridoxine dependent epilepsy

**DOI:** 10.1101/2025.05.07.652392

**Authors:** Imke M.E. Schuurmans, Udo Engelke, Muna Abedrabbo, Sofía Puvogel, Rachel Mijdam, Gijs-Jan Scholten, Sara B. van Katwijk, Astrid Oudakker, Hilal H. Al-Shekaili, Dirk J. Lefeber, Blair R. Leavitt, Clara D.M. van Karnebeek, Nael Nadif Kasri, Alejandro Garanto

## Abstract

Pyridoxine-dependent epilepsy (PDE) is a rare neurometabolic disorder of lysine catabolism caused by bi-allelic variants in *ALDH7A1*. This enzyme deficiency leads to the accumulation of neurotoxic metabolites, pyridoxal-phosphate inactivation and consequently severe neurological symptoms. Current treatments, including vitamin B6 supplementation and lysine-restricted diets, partially alleviate seizures and intellectual disability but are not curative. To explore underlying mechanisms and potential therapies, we generated patient-derived human induced pluripotent stem lines (hiPSC) that were subsequently differentiated into astrocytes, the primary source of *ALDH7A1* in the brain and key regulators of metabolic homeostasis. Metabolomic analyses confirmed elevated PDE biomarkers, and RNA sequencing revealed gene expression changes consistent with increased oxidative stress. Oxidative damage was validated by markers of DNA oxidation and lipid peroxidation. In addition, dysregulated oxygen consumption rates suggested mitochondrial dysfunction in PDE astrocytes. Notably, these pathological phenotypes were alleviated by downregulating *AASS*, the first enzyme of the lysine catabolism, by using CRISPR/Cas9 editing or antisense oligonucleotides (AON). This demonstrates that lysine catabolism underlies these phenotypes and highlights the therapeutic potential of AON therapy targeting *AASS* to reduce neurotoxic metabolite accumulation. These findings provide a promising strategy for developing targeted treatments for PDE and other rare neurometabolic disorders.

## Introduction

Pyridoxine-dependent epilepsy (PDE) is a rare neurometabolic disorder of lysine catabolism, with an estimated incidence around 1:65,000^1^. PDE is characterized by recurrent perinatal-onset seizures that are resistant to conventional anticonvulsant drugs but instead show remarkable response to pyridoxine administration^2^. In addition to epilepsy, more than 75% of the PDE patients exhibit developmental delay and moderate to severe intellectual disability^3^, in some cases combined with structural brain abnormalities including hypoplasia of the corpus callosum and/or cerebellum^4,5^. PDE is caused by autosomal recessive pathogenic variants in *ALDH7A1*. The encoded enzyme is responsible for the conversion of 2-aminoadipic-6-semialdehyde (α-AASA) to 2-aminoadipic acid (AAA) in the lysine degradation pathway. ALDH7A1 enzyme deficiency therefore results in accumulation of neurotoxic metabolites including α-AASA, 1-piperideine-6-carboxylic acid (P6C) and pipecolic acid (PA). Excessive P6C is known to complex with pyridoxal phosphate (PLP), resulting in its lowered availability and therefore PLP inactivation^1^. PLP is the active form of vitamin B6 (pyridoxine) acting as an important cofactor for several enzymes in the brain, including enzymes involved in GABA synthesis^6^. PLP inactivation is therefore thought to underly the pyridoxine-responsive seizures. Astrocytes are the main cell type expressing *ALDH7A1* in the brain^7^. Since the accumulation of toxic lysine degradation intermediates primarily occurs in these cells they are thought to play a crucial role in the disease mechanisms underlying PDE^2,7–11^, but the exact mechanisms remain unclear.

Currently, the only available treatment strategies for PDE include vitamin B6 (pyridoxine) supplementation for seizure control combined with dietary interventions aimed at reducing toxic metabolite accumulation by lysine reduction and arginine supplementation. Arginine is included as a competitive inhibitor for lysine to cross the blood-brain barrier as lysine and arginine use the same cationic transporter^12^. However, these dietary interventions have limited efficacy as lysine is an essential amino acid that cannot be completely removed. Thus, there is an unmet need to develop novel therapeutic strategies as lysine reduction does not eliminate, but at best mitigates neurologic impairments^13^.

Upstream enzyme inhibition to prevent buildup of toxic metabolites has been successful for other inherited metabolic disorders (IMDs) such as tyrosinemia type I^14^. A similar strategy could provide an alternative therapy for PDE instead of current dietary interventions. AASS is a bifunctional enzyme, consisting of the lysine ketoglutarate reductase (LKR) domain and saccharopine dehydrogenase (SDH) domain, catalyzing the first and second step respectively of the saccharopine pathway of lysine catabolism. Importantly, near-complete AASS deficiency in humans, presents with a benign phenotype causing no or mild symptoms only^15^. This provides most direct evidence that AASS could be a safe target for therapeutic inhibition. Several preclinical studies already showed the effectiveness of inhibiting AASS to treat glutaric aciduria type I (GA1), another disorder of lysine catabolism^16,17^. Given that AASS works upstream of ALDH7A1 in the lysine pathway, we propose AASS as a potential target for therapeutic inhibition to prevent the cerebral accumulation of toxic lysine metabolites in PDE. A promising strategy for partial inhibition of AASS are antisense oligonucleotides (AONs). These are small nucleic acid molecules that bind complementarily to the pre-mRNA or mRNA and thereby, among other functions, can modulate degradation of mRNA transcripts^18^. Several AON strategies have already demonstrated promising results in clinical trials for other inherited diseases, particularly those affecting the eye or brain, due to the benefits of local delivery^19^.

Currently available PDE model systems include *Aldh7a1*^-/-^ mice and zebrafish, both recapitulating some of the typical PDE characteristics, including the pyridoxine-dependent seizures^2,20^. Although these PDE animal models have improved our understanding of the PDE disease mechanisms, the use of animal models also has limitations including functional and genomic differences^21,22^. Human induced pluripotent stem cells (hiPSCs) offer opportunities for research into human and cell-type-specific pathophysiological mechanisms underlying PDE and allow us to test new genetic therapeutic strategies for PDE in a patient’s context.

To this end, we developed an isogenic *ALDH7A1*^-/-^ (*ALDH7A1* KO) hiPSC line and generated three PDE patient-derived hiPSC-lines. Considering the enriched expression of *ALDH7A1* in astrocytes compared to other neuronal cell types and their previously suggested role in the pathophysiology of PDE, we differentiated the *ALDH7A1* KO and PDE patient-derived hiPSC lines into astrocytes. Our data show that the *ALDH7A1* KO as well as the PDE patient astrocytes show elevated PDE biomarkers, increased oxidative stress and heightened oxygen consumption rates (OCRs), indicative of mitochondrial dysfunction. Notably, we demonstrated that partial inhibition of *AASS*, the first enzyme in lysine catabolism, using AONs effectively ameliorated these metabolic abnormalities and normalized OCRs.

## Results

### PDE patient-derived astrocytes show elevated PDE biomarkers

To investigate the disease mechanism underlying PDE, we generated hiPSC lines from PDE patients^23^ and an isogenic hiPSC *ALDH7A1* knock-out line^24^. Fibroblasts from three PDE patients (P1-P3) were reprogrammed towards hiPSCs, all harbouring the most commonly reported *ALDH7A1* homozygous c.1279G>C (p.Glu427Gln) variant (Fig. 1A). We used CRISPR/Cas9 genome editing in a healthy control line (C1) to generate an isogenic *ALDH7A1* KO line (Fig. 1B). As expected, western blotting revealed full loss of ALDH7A1 protein levels in the isogenic *ALDH7A1* KO line^24^. To further minimalize potential effects of genetic background, we also included four independent control lines (C1-C4) for comparison.

**Figure 1.**
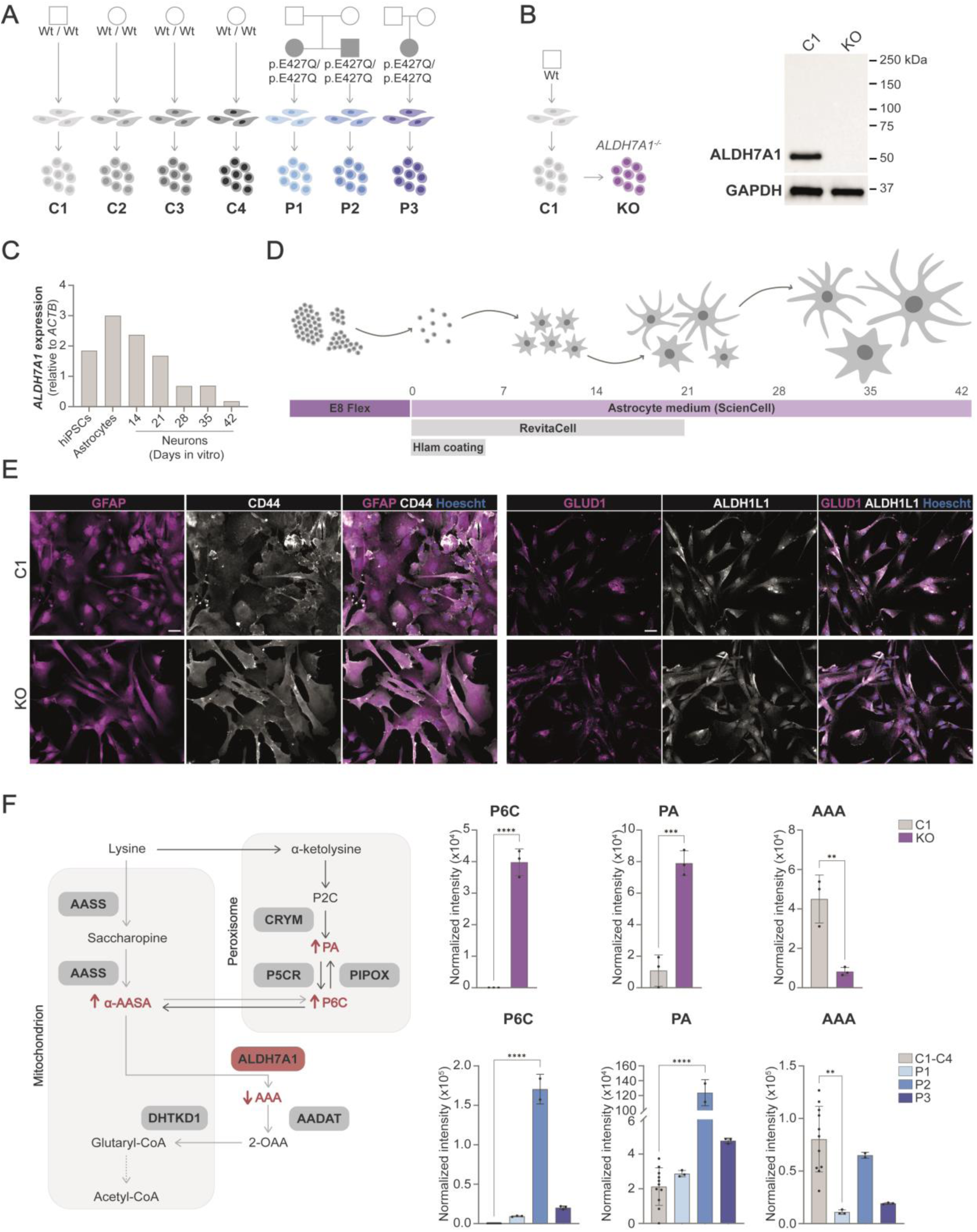
Metabolic characterization of *ALDH7A1* KO and PDE patient-derived hiPSC-astrocytes. **A.** Schematic overview of the four control and three PDE-patient hiPSC lines used in this study carrying the pathogenic *ALDH7A1* variant c.1279G>C (p.Glu427Gln) in homozygosis. **B.** Schematic overview of *ALDH7A1* KO (KO) hiPSC line and C1 including previously published western blot for ALDH7A1 and GAPDH protein levels for the KO and its control line^24^.**C.** Expression of *ALDH7A1* relative to *ACTB* by RT-PCR in hiPSCs, hiPSC-derived astrocytes and hiPSC-derived neurons from 14, 21, 28, 35 and 42 days *in vitro* (DIV). **D.** Schematic representation of the protocol to differentiate hiPSCs towards astrocytes. **E.** Representative images of immunostaining of GFAP (magenta), CD44 (white), GLUD1 (magenta) and ALDH1L1 (white) in DIV 35 C1 and KO astrocytes. All pictures were taken at the same magnification (scale bar = 50 µm). **F.** Schematic representation of ALDH7A1 deficiency in the lysine pathway including the corresponding PDE biomarkers. The normalized intensity (peak area) of P6C, PA and AAA measured via NGMS in KO and C1 astrocytes is shown in the top three graphs. *n* = 3 for C1; *n* = 3 for KO. Statistically significant differences were tested through unpaired t-test. In the bottom three graphs the normalized intensity of the same biomarkers are shown for the three PDE patient lines as well as the four control astrocyte lines (C1-C4). *n* = 12 for C1-C4; *n* = 3 for P1; *n* = 2 for P2; *n* = 3 for P3. Statistically significant differences for P6C and PA were tested through one-way ANOVA and Dunnett’s multiple comparison test, while Kruskal-Wallis test combined with Dunn’s multiple comparison test was used for the AAA biomarker. For all graphs in this figure statistical significance is indicated by: \**P* < 0.05, \*\**P* < 0.01, \*\*\**P* < 0.005, \*\*\*\**P* < 0.0001. Exact *P-*value per condition is provided in Supplementary Table 1.

Astrocytes are thought to play a key role in the pathophysiology of PDE^2,7,10^, especially as the expression of *ALDH7A1* is enriched in astrocytes compared to neurons^7^. We confirmed that *ALDH7A1* expression was higher in hiPSC-astrocytes compared to hiPSCs and hiPSC-neurons, in which the expression of *ALDH7A1* rapidly decreases during differentiation (Fig. 1C and Supplementary Fig. 1A). Given these findings, we considered hiPSC-derived astrocytes to be most suitable for further studying PDE disease mechanisms. We differentiated the hiPSC-lines towards astrocytes (Fig. 1D), as described recently^25^. All hiPSC-lines differentiated towards astrocytes, without observable differences between cell lines. At days in vitro (DIV) 35, the astrocytes showed expression of typical astrocyte markers including GFAP, CD44, ALDH1L1 and GLUD1 (Fig. 1E and Supplementary Fig. 1B). In addition, the hiPSC-derived astrocyte cultures showed typical monoculture astrocyte morphology.

To investigate the metabolic phenotype of the PDE astrocytes we measured several intracellular PDE biomarkers, including P6C, PA and AAA, under basal conditions using next-generation metabolic screening (NGMS, Fig. 1F and Supplementary Fig. 2). As expected, P6C and PA were significantly increased in the *ALDH7A1* KO astrocytes compared to control, while AAA was significantly decreased. A similar overall metabolic profile was observed in PDE patient astrocytes, though with greater variability between individual lines. P6C levels were significantly elevated across all PDE patient lines compared to controls, in which P6C remained undetected. Additionally, PA levels were significantly higher in P2. Whereas AAA levels showed considerable variability in the control group, AAA levels were significantly lower in P1 astrocytes. Overall, the metabolic profiles of *ALDH7A1* KO and PDE patient astrocytes align with the metabolic phenotypes observed in PDE patients *in vivo*.

### Increased oxidative stress and impaired oxygen response in PDE astrocytes

To explore for the disease mechanisms underlying PDE, we analyzed transcriptional differences in PDE astrocytes using bulk RNA-sequencing of two controls (C1 and C4), two PDE patient lines (P1 and P3) and *ALDH7A1* KO astrocytes. Principal component (PC) analysis confirmed transcriptional changes across hiPSC-astrocyte cultures, clearly separating the different lines (Supplementary Fig. 3). Subsequent differential gene expression analysis was conducted independently for the *ALDH7A1* KO astrocytes with respect to their isogenic control (Fig. 2A), and for the PDE patient astrocytes in comparison to pooled controls (Fig. 2B). This analysis revealed a total of 4811 down-regulated and 5170 up-regulated genes (Fig. 2C) in the *ALDH7A1* KO astrocytes compared to C1 astrocytes. The number of differentially expressed genes (DEGs) in PDE patient astrocytes versus controls was lower, with 355 down-regulated and 375 up-regulated genes (Fig. 2C). Among these transcriptomic profiles, we sought to identify pathways most relevant to PDE. We performed gene ontology (GO) enrichment analysis in both the isogenic pair and PDE patient astrocytes, focusing on shared pathways affected across all PDE astrocyte lines (Fig. 2D). Upregulated pathways were involved in ion binding, development, migration, and growth factors, while downregulated pathways were primarily associated with cell signaling, particularly oxygen response, indicating a potential increase in oxidative stress in PDE astrocytes, as has been previously proposed^2,11,26^. The ALDH7A1 enzyme has also been described to protect against hyperosmotic stress through the generation of an important cellular osmolyte, formed from betaine aldehyde^27,28^. Hyperosmotic stress is coupled to an increase in oxidative stress through generation of reactive oxygen species (ROS) as well as lipid peroxidation (LPO)^29^. Additionally, ALDH7A1 has been identified to remove several LPO-derived aldehydes which are formed under oxidative conditions, which in turn leads to increased affinity of ALDH7A1 for these toxic aldehydes^27,28^. The strong correlation between osmotic and oxidative stress indicates that cytoprotective roles of ALDH7A1 may be twofold: (1) producing osmolytes to mitigate osmotic stress and (2) eliminating reactive aldehydes formed due to increased oxidative stress.

**Figure 2.**
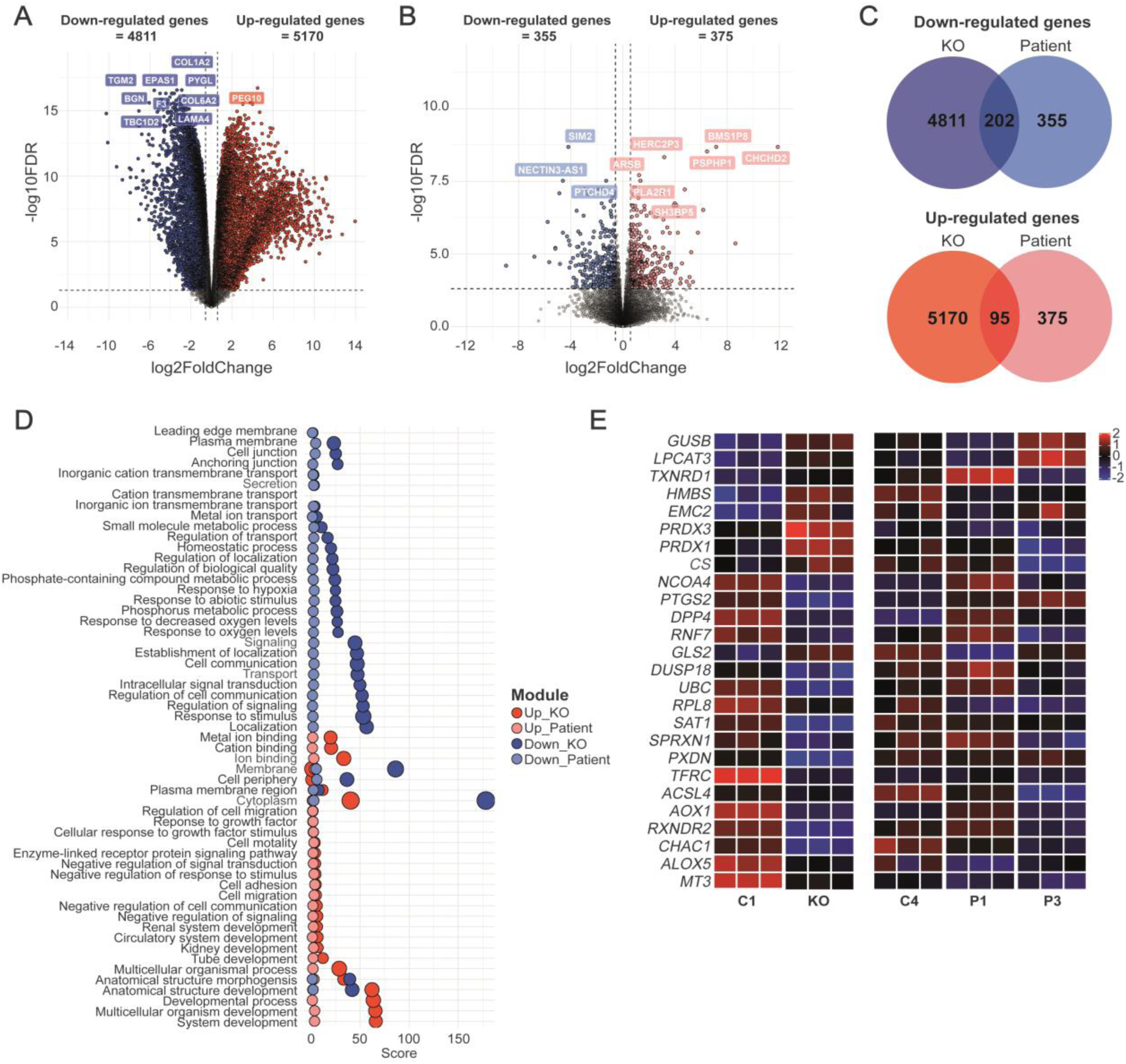
RNA sequencing in PDE patient-derived astrocytes suggests impaired oxygen response and increased oxidative stress. **A.** Volcano plot depicting differentially expressed genes (DEGs) between *ALDH7A1* KO (KO) and C1 astrocytes. Genes were defined as differentially expressed when the absolute log_2_ fold change (Log_2_FC) exceeds 0.58 and with a Benjamini-Hochberg (BH)-adjusted p-value below 0.05. Colored dots indicate DEGs; down-regulated genes in KO astrocytes are depicted in blue, while up-regulated genes are shown in red. The ten genes with the largest difference in expression between KO and C1 astrocytes are indicated. **B.** Volcano plot depicting DEGs between astrocytes from P1 and P3 compared to C1 and C4. Genes were defined as differentially expressed when the absolute log_2_ fold change (Log_2_FC) exceeds 0.58 and with a BH-adjusted *p*-value below 0.05. Colored dots indicate DEGs; down-regulated genes in KO astrocytes are depicted in light-blue, while up-regulated genes are shown in light-red. The ten genes with the largest difference in expression between PDE and control astrocytes are indicated. **C.** Venn diagram depicting the number of shared up-regulated genes (red) and down-regulated genes (blue) between the list of DEGs in KO versus C1 astrocytes and between astrocytes from P1 and P3 versus to C1 and C4. **D.** Scatterplot showing the enriched GO terms of the shared genes per ontological category, associated with up-regulated (red dots) and down-regulated genes (blue dots) of both the DEGs between the KO versus C1 astrocytes as well as the between the P1 and P3 astrocytes versus C1 and C4 astrocytes. The size of the dots indicates the number of intersected genes between the list of DEGs and the genes associated with the particular term. **E.** Heatmap of gene expression from the C1, KO, C4, P1 and P3 astrocytes of genes associated with lipid peroxidation related oxidative stress.

Considering the potential role of ALDH7A1 in oxidative stress, we examined transcriptional differences in LPO-related genes in PDE astrocytes (Fig. 2E). *LPCAT3* upregulation suggests changes in lipid metabolism that heighten lipid peroxidation vulnerability^30^. Increased expression of *TXNRD1*, *PRDX3*, and *PRDX1* indicates a response to elevated LPO-related ROS^31–33^, while *PXDN* downregulation hampers oxidative stress mitigation^34^. Downregulation of *ACSL4*, *ALOX5*, and *CHAC1* impacts lipid metabolism and glutathione defenses, reducing the ability to counteract oxidative stress^35–37^. These changes imply a decreased capacity to manage oxidative stress and a higher risk of LPO damage.

To directly probe for oxidative stress in PDE astrocytes we measured 8-Oxo-dG under baseline conditions using immunocytochemistry. 8-Oxo-dG is one of the major DNA oxidation products and therefore generally considered a measure for oxidative stress. We observed a significant increased signal for 8-Oxo-dG in the *ALDH7A1* KO astrocytes compared to C1 (Fig. 3A). As ALDH7A1 has been predicted to play a role in lipid metabolism, we also measured LPO-related oxidative stress through 4-HNE immunostaining, one of the main metabolites produced during LPO. We observed the 4-HNE fluorescent signal to be significantly increased in the *ALDH7A1* KO astrocytes compared to C1 (Fig. 3B). In contrast to the *ALDH7A1* KO astrocytes, we did not observe an increase of the 8-Oxo-dG signal (Fig. 3C) in any of the PDE patient astrocyte lines. The 4-HNE signal (Fig. 3D) was however significantly elevated in P1 and P3 astrocytes compared to control (C1-C4) astrocytes. Overall, PDE astrocytes exhibit elevated signs of oxidative stress levels; however, these levels are more pronounced in *ALDH7A1* KO astrocytes compared to PDE patient astrocytes.

**Figure 3.**
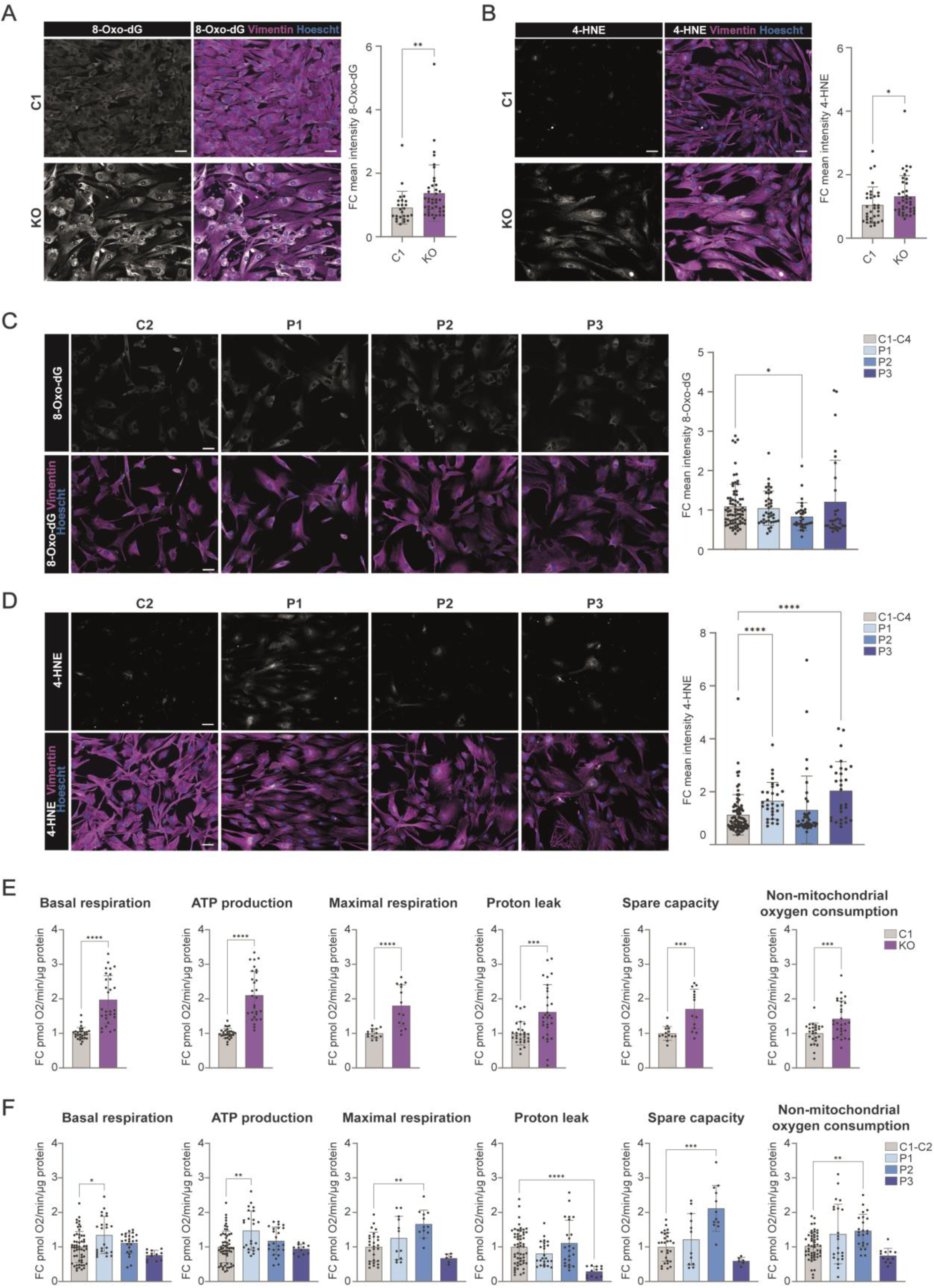
Increased oxygen consumption rates and oxidative stress levels in PDE astrocytes. **A.** Representative images of 8-Oxo-dG immunostaining and FC of mean intensity of 8-Oxo-dG per well relative to average intensity of C1 shown for *ALDH7A1* KO (KO) and C1 astrocytes. *n* = 25/6 for C1; *n* = 40/6 for KO **B.** Representative images of 4-HNE immunostaining and FC of mean intensity of 4-HNE per well relative to average intensity of C1 shown for KO and C1 astrocytes. *n* = 33/4 for C1; *n* = 39/4 for KO. For **A.** and **B.**, statistically significant differences were tested through Mann-Whitney test. **C.** Representative images of 8-Oxo-dG immunostaining and FC of mean intensity of 8-Oxo-dG per well relative to average intensity of merged controls shown for P1, P2, P3 and merged controls (C1-C4). *n* = 75/6 for C1-C4; *n* = 40/6 for P1; *n* = 32/5 for P2; *n* = 30/6 for P3. **D.** Representative images of 4-HNE immunostaining and FC of mean intensity of 4-HNE per well relative to average intensity of merged controls shown for P1, P2, P3 and merged controls (C1-C4). *n* = 66/4 for C1-C4; *n* = 31/4 for P1; *n* = 39/4 for P2; *n* = 29/4 for P3.For **C.** and **D.**, statistically significant differences were tested through Kruskal-Wallis test combined with Dunn’s testing. **E.** Fold Change (FC) of Basal respiration (BR), ATP production (AP), maximal respiration (MR), proton leak (PL), spare capacity (SC) and non-mitochondrial oxygen consumption (NMOC) is depicted for KO and C1 astrocytes at DIV 35, as measured with seahorse assay. For BR, AP, PL and NMOC: *n* = 28/2 for C1; *n* = 29/2 for KO. For MR and CP: *n* = 14/2 for C1; *n* = 14/2 for KO. Statistically significant differences were tested through unpaired t test or Mann-Whitney test. **F.** FC of BR, AP, MR, PL, SC and NMOC is depicted for DIV 35 astrocytes from P1, P2, P3 and merged controls (C1 and C2). For BR, AP, PL and NMOC: *n* = 56/2 for C1-C4; *n* = 24/2 for P1; *n* = 24/2 for P2; *n* = 12/1 for P3. For MR and CP: *n* = 28/2 for C1-C4; *n* = 12/2 for P1; *n* = 12/2 for P2; *n* = 6/1 for P3.Statistically significant differences were tested through ordinary one-way ANOVA and Dunnett’s multiple comparison test or with Kruskal-Wallis test combined with Dunn’s multiple comparison test. For all graphs in this figure statistical significance is indicated by: *P < 0.05, **P < 0.01, ***P < 0.005, ****P < 0.0001. Scale bar = 50 μm. Exact *P-*value per condition is provided in Supplementary Table 1.

The lysine degradation pathway is partially localized within the mitochondria, so the accumulation of toxic lysine intermediates is thought to contribute directly to mitochondrial dysfunction^38–40^. Furthermore, increased ROS and the resulting oxidative stress are also known to impair mitochondrial function^41–44^. Given these facts and the elevated oxidative stress identified in PDE astrocytes, we further investigated mitochondrial function in these cells. We performed Seahorse assay to measure cellular respiration by monitoring the OCR. We included astrocytes derived from the three patients, the *ALDH7A1* KO and its control (C1), and an additional control line (C2). We observed a significant increase for all OCR parameters in *ALDH7A1* KO astrocytes (Fig. 3E and Supplementary Fig. 4), suggesting a compensatory upregulation of mitochondrial activity, likely as an adaptive response to the accumulation of toxic lysine intermediates and the associated metabolic and oxidative stress. Similarly, some of these parameters were significantly elevated in P1 and P2 astrocytes (Fig. 3F and Supplementary Fig. 4), but not in P3 astrocytes, which instead showed a reduction in proton leak compared to control. Overall, PDE astrocytes exhibited an increased OCR, but in line with oxidative stress measurements, this increase was more pronounced in *ALDH7A1* KO astrocytes than in PDE patient astrocytes.

### Rescue of the metabolic and cellular phenotypes in *ALDH7A1* KO astrocytes by targeting the upstream enzyme AASS

Considering that the accumulation of the toxic lysine derivatives in PDE are causative for the neurologic phenotype, therapeutic strategies for PDE should aim to reduce this accumulation by blocking the entrance of lysine into the catabolic pathway. As previously indicated, increased lysine levels due to deficiency in the first step of the lysine pathway catalyzed by AASS have not caused any relevant clinical phenotype^15^. Moreover, *AASS* inhibition to treat other lysine metabolism disorders such as GA1 has already been described^16,17^. Therefore, we considered *AASS* a potential target for therapeutic inhibition to reduce the accumulation of toxic lysine derivatives in PDE. To test this hypothesis, we first generated a double knock-out (DKO) hiPSC line for *ALDH7A1* and *AASS* (*ALDH7A1/AASS* DKO) by targeting *AASS* through CRISPR/Cas9 genome editing in the *ALDH7A1* KO hiPSC line (Supplementary Fig. 5). Western blotting of the *ALDH7A1* KO hiPSC line and *ALDH7A1/AASS* DKO hiPSC line confirmed full knock-out of ALDH7A1 in both lines, and around 95% reduction of AASS levels in the *ALDH7A1/AASS* DKO hiPSC line compared to C1 (Fig. 4A). Notably, we observed that AASS levels were already reduced by approximately 50% in the *ALDH7A1* KO hiPSC line compared to C1, suggesting that cells might already downregulate *AASS* as homeostatic response to loss of *ALDH7A1*.

**Figure 4.**
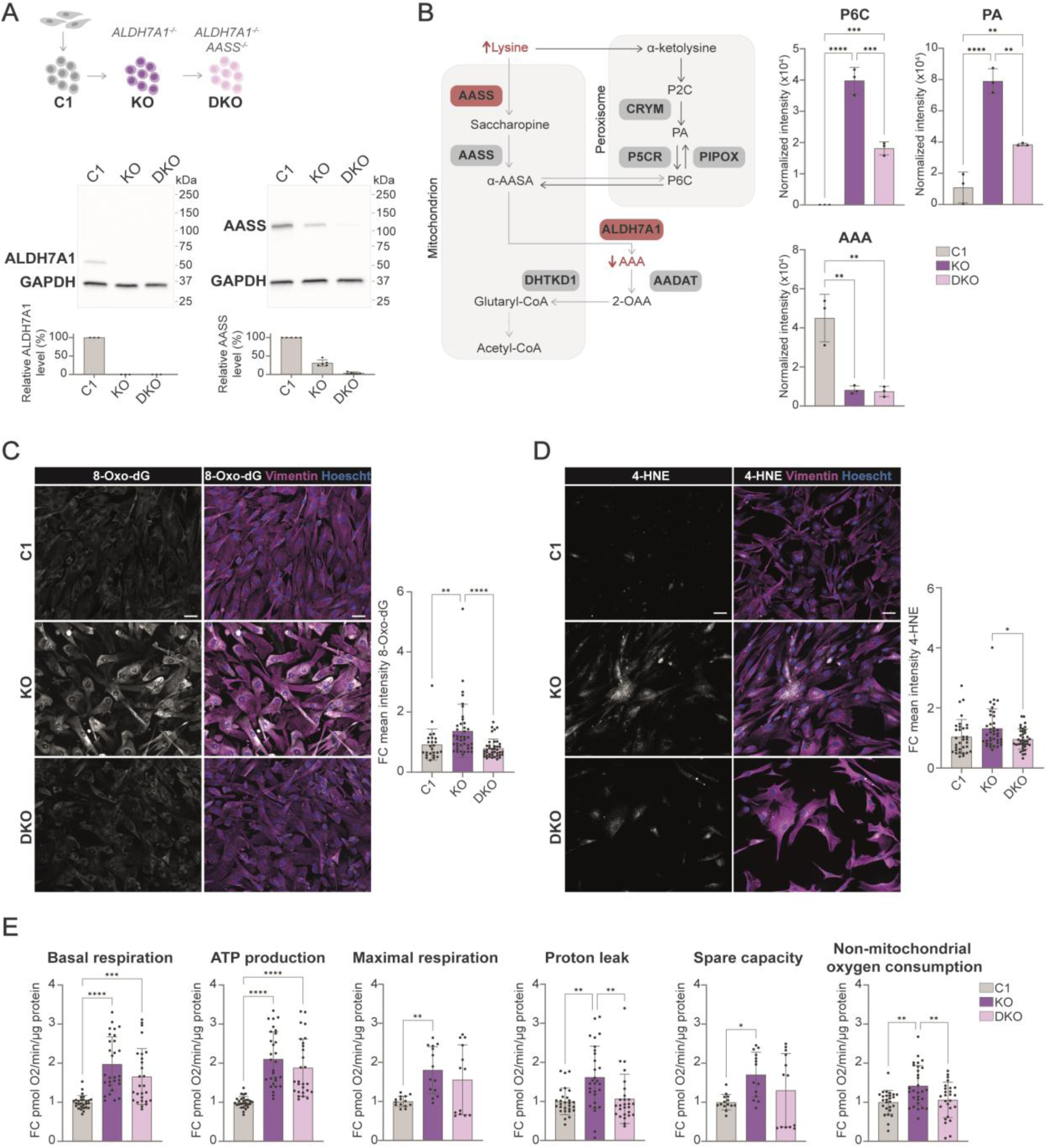
Rescue of metabolic and cellular phenotypes in PDE astrocytes by the decrease of AASS enzyme. **A.** Schematic overview of *ALDH7A1* KO (KO) hiPSC line and *ALDH7A1/AASS* DKO (DKO) hiPSC line, including western blot for AASS, ALDH7A1 and GAPDH proteins. **B.** Schematic representation of ALDH7A1 deficiency combined with AASS deficiency in the lysine pathway including the corresponding biomarkers. The normalized intensity of P6C, PA and AAA measured via NGMS in KO, DKO and C1 astrocytes is depicted. *n* = 3 for C1; *n* = 3 for KO; *n* = 3 for DKO. Statistically significant differences were tested through ordinary one-way ANOVA and Tukey’s multiple comparison test. **C.** Representative images of 8-Oxo-dG immunostaining (Scale bar = 50 μm) and FC of mean intensity of 8-Oxo-dG per well relative to average intensity of C1 shown for KO, DKO and C1 astrocytes. *n* = 25/6 for C1; *n* = 40/6 for KO; *n* = 40/6 for DKO. **D.** Representative images of 4-HNE immunostaining (Scale bar = 50 μm) and FC of mean intensity of 4-HNE per well relative to average intensity of C1 shown for KO, DKO and C1 astrocytes. *n* = 34/4 for C1; *n* = 39/4 for KO; *n* = 38/4 for DKO. For both **C.** and **D.**, statistically significant differences were tested through Kruskal-Wallis test combined with Dunn’s testing. **E.** Fold change (FC) of Basal respiration (BR), ATP production (AP), proton leak (PL), maximal respiration (MR), spare capacity (SC) and non-mitochondrial oxygen consumption (NMOC) is depicted for KO, DKO and C1 astrocytes at DIV 35. For BR, AP, PL and NMOC: *n* = 28/2 for C1; *n* = 29/2 for KO; *n* = 27/2 for DKO. For MR and CP: *n* = 14/2 for C1; *n* = 14/2 for KO; *n* = 13/2 for DKO. Statistically significant differences were tested through unpaired t test or Mann-Whitney test. For all graphs in this figure statistical significance is indicated by: *P < 0.05, **P < 0.01, ***P < 0.005, ****P < 0.0001. Exact *P-*value per condition is provided in Supplementary Table 1.

The *ALDH7A1* KO, *ALDH7A1/AASS* DKO and C1 lines were differentiated towards astrocytes, all showing expression of the astrocyte markers and no effect on differentiation due to additional KO of *AASS* (Fig. 1E and Supplementary Fig. 1B). To investigate whether targeting *AASS* in *ALDH7A1* KO astrocytes could rescue the metabolic phenotype, we measured the PDE biomarkers under basal conditions using NGMS (Fig. 4B and Supplementary Fig. 2). As expected, AAA levels remained low in the *ALDH7A1/AASS* DKO astrocytes, similar to *ALDH7A1* KO astrocytes. P6C and PA levels in the *ALDH7A1/AASS* DKO astrocytes were significantly reduced, but did not reach control levels. This suggests that while targeting *AASS* in *ALDH7A1* KO astrocytes mitigates the metabolic imbalance, it does not fully rescue it. We next assessed whether this partial metabolic restoration was sufficient to reduce oxidative stress in *ALDH7A1* KO astrocytes. In the *ALDH7A1/AASS* DKO astrocytes both 8-Oxo-dG (Fig. 4C) and 4-HNE (Fig. 4D) levels were fully normalized to control levels. This suggests a rescued oxidative stress response upon partial metabolic rescue by targeting *AASS* in *ALDH7A1* KO astrocytes.

Finally, we investigated if *AASS* inhibition could also restore the increased OCR observed in the *ALDH7A1* KO astrocytes (Fig. 4E). We observed that the proton leak and non-mitochondrial oxygen consumption were fully restored in the *ALDH7A1/AASS* DKO astrocytes. However, the basal respiration, ATP production, maximal respiration and spare capacity were not completely normalized. Altogether, these findings indicate that by targeting *AASS* in PDE astrocytes both the metabolic phenotype as well as cellular phenotypes, defined by increased oxidative stress and OCR, can be ameliorated.

### AON-mediated downregulation of *AASS* can partially rescue metabolic and cellular phenotypes in PDE patient astrocytes

To recreate a situation comparable to PDE-affected individuals, in which *AASS* is present during development and therefore can only be targeted after development, we designed an alternative strategy in which we partially reduced *AASS* levels using antisense technology. Two distinct AON strategies, splice-switching AONs (ssAONs) and gapmers, were employed to target and promote the degradation of *AASS* transcripts (Supplementary Fig. 6A). ssAONs (S1, S2 and S3) were designed to target a splice site (enhancer) to promote exon skipping, resulting in an out-of-frame transcript and subsequently degradation by nonsense-mediated decay. In contrast, gapmers (G1, G2 and G3; Supplementary Fig. 6B) contain a DNA core sequence flanked by RNA wings that binds to the pre- and mRNA to degrade the transcript through RNase H recruitment. Our AONs were designed to target the region encoding for the LKR domain of *AASS* (the large *AASS* isoform) leaving the SDH domain (shortest isoform) intact. We specifically aimed to target the LKR AASS activity, as the accumulation of saccharopine (the product generated by the SDH activity) has been shown to be neurotoxic, while the accumulation of lysine seems to be safe^45^.

To evaluate the efficiency of our AONs to downregulate *AASS*, wildtype fibroblasts were transfected with liposomes including the ssAONs (0.5 or 1.0 µM), gapmers (0.2 or 0.5 µM) or the respective negative sense control (Son) corresponding to the ssAONs (SonS; at 1.0 µM) or gapmers (SonG; at 0.5 µM). Four days post-delivery *AASS* downregulation was assessed at RNA level through RT-qPCR (Supplementary Fig.6C) and protein level (Supplementary Fig. 6D) by western blot. For the RT-qPCR, amplicons spanning the region between exons 4-5 and exons 6-8 of the *AASS* transcript were amplified, as these exons are unique to the large *AASS* isoform and specifically encode the LKR domain. Although we observed efficient exon skipping upon transfection with the ssAONs, this did not result in degradation of the *AASS* transcript (Supplementary Fig. 6C and 6D). We therefore excluded the ssAONs from following experiments. Transfection with the gapmers resulted in approximately 50-80% downregulation of AASS levels relative to SonG control (Supplementary Fig. 6C and 6D). We selected G3 for further validation in astrocytes. Accordingly, four- to five-week-old C1 astrocytes were transfected for seven days with G3 using liposomes at concentrations ranging from 0.05 - 0.5 µM. We observed approximately 50% remaining *AASS* expression with RT-qPCR (Supplementary Fig. 7A) and around 10% remaining AASS levels with western blotting (Supplementary Fig. 7B) upon G3 delivery at 0.5 µM in C1 astrocytes. Finally, at 0.5 µM G3 resulted in significant reduction of *AASS* expression relative to SonG in both the *ALDH7A1* KO and PDE patient astrocytes, ranging from 50-70% downregulation (Fig. 5A and Supplementary Fig. 7C). *AASS* downregulation of G3 in P2 astrocytes showed ∼50% reduction of AASS levels (Fig. 5B).

**Figure 5.**
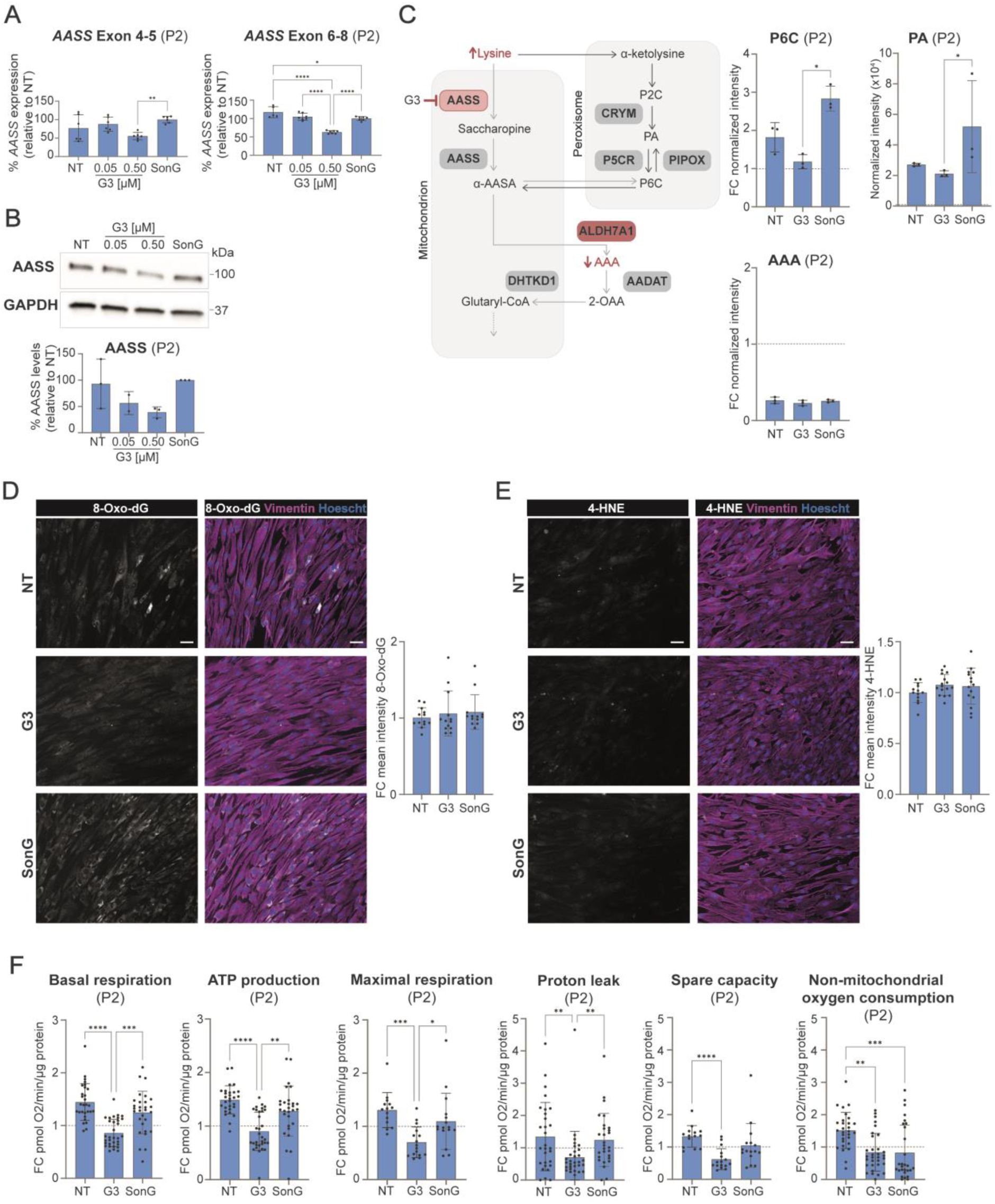
Gapmer-Mediated Rescue of Metabolic Phenotype and Oxygen Consumption in P2 Astrocytes. All experiments included non-treated control (NT), G3 at 0.5 µM (and 0.05 µM), and a sense oligonucleotide (SonG) control at 0.5 µM. **A.** Relative expression of *AASS* exon 4-5 and exon 6-8 in P2 astrocytes seven days post-G3 delivery, assessed by qPCR and normalized to *GUSB*. Expression is shown as a percentage relative to SonG. For *AASS* exon 4-5: *n* = 6/2 for NT; *n* = 6/2 for G3 0.05 µM; *n* = 6/2 for G3 0.5 µM; *n* = 6/2 for SonG. For *AASS* exon 6-8: *n* = 5/3 for NT; *n* = 7/3 for G3 0.05 µM; *n* = 7/3 for G3 0.5 µM; *n* = 7/3 for SonG. Statistical significance was determined by ordinary one-way ANOVA with Dunnett’s multiple comparison test. **B.** Semi-quantification of AASS protein levels relative to GAPDH, with representative (cropped) western blot image (*n* = 3) of P2 astrocytes seven days post-G3 delivery. Statistical significance was tested using one-way ANOVA with Dunnett’s multiple comparison test. **C.** Schematic of ALDH7A1 deficiency and G3-mediated *AASS* downregulation in the lysine pathway, highlighting corresponding biomarkers. Normalized PA intensity and fold change (FC) of P6C and AAA levels via NGMS are presented for NT, G3, and SonG conditions. *n* = 3 for NT; *n* = 3 for G3; *n* = 3 for SonG. Statistical significance was assessed via one-way ANOVA and Tukey’s multiple comparison test. **D-E.** Representative immunostaining images (scale bar = 50 μm) and FC of mean intensity per well relative to NT for 8-Oxo-dG **(D)** and 4-HNE **(E)** under NT, G3, and SonG conditions. For 8-Oxo-dG immunostaining: *n* = 13/2 for NT; *n* = 13/2 for G3; *n* = 13/2 for SonG. For 4-HNE immunostaining: *n* = 13/2 for NT; *n* = 16/2 for G3; *n* = 15/2 for SonG. Statistical significance was determined via Kruskal-Wallis test with Dunn’s post hoc test. **F.** FC values of basal respiration (BR), ATP production (AP), proton leak (PL), maximal respiration (MR), spare capacity (SC), and non-mitochondrial oxygen consumption (NMOC) for NT, G3, and SonG conditions in P2 astrocytes. For BR, AP, PL and NMOC: *n* = 29/2 for NT; *n* = 33/2 for G3; *n* = 29/2 for SonG. For MR and CP: *n* = 14/2 for NT; *n* = 16/2 for G3; *n* = 16/2 for SonG. Statistical significance was tested using unpaired t-test or Mann-Whitney test. For all graphs, statistical significance is indicated as *P < 0.05, **P < 0.01, ***P < 0.005, ****P < 0.0001. Exact *P-*value per condition is provided in Supplementary Table 1.

We then assessed whether 50% downregulation of *AASS* in PDE patients could mitigate the metabolic and cellular phenotypes. As the metabolic and OCR parameters were most severely affected in the P2 astrocytes, we used this patient line for the functional validations. The metabolic phenotype was assessed using NGMS seven days upon G3-delivery (0.5 µM) including non-treated (NT) and SonG control (Fig. 5C). As expected, AAA levels were unchanged in the G3-treated P2 astrocytes but still lower compared to control (indicated by the dotted line). Notably, P6C levels were almost normalized to control levels in the G3-treated P2 astrocytes. Although PA levels were improved in the G3-treated condition of the P2 astrocytes, it was not fully normalized to control levels.

We next evaluated oxidative stress levels following treatment with G3 (0.5 µM) for seven days. Consistent with previous findings, immunostaining for 8-Oxo-dG and 4-HNE did not reveal increased levels of these markers compared to control at baseline (Supplementary Fig. 8A-B), and no significant difference was observed between the G3 and control conditions (Fig. 5D-E and Supplementary Fig. 8C-D). This indicates that when no baseline differences are present, *AASS* inhibition does not seem to affect oxidative stress levels. However, increased levels of 8-Oxo-dG and 4-HNE in P1 and P3 (Supplementary Fig. 8) were also not rescued upon G3 treatment.

Lastly, we evaluated the ability of G3 (0.5 µM) to reduce the OCR in P2 astrocytes (Fig. 5F). We observed that the basal respiration, ATP production and non-mitochondrial oxygen consumption were completely normalized to control levels in the P2 astrocytes upon G3 treatment. The proton leak, maximal respiration and spare capacity were also significantly reduced in the G3-condition compared to the NT condition, but reached even lower levels compared to control. The non-mitochondrial oxygen consumption was also significantly reduced in the SonG condition compared to NT, of which the effects and/or toxicity could be either chemistry or sequence dependent^46^. Altogether, our data indicate that G3-mediated downregulation of *AASS* in mature astrocytes is able to partially restore PDE biomarkers and increased OCR.

## Discussion

In this study, we used isogenic *ALDH7A1* KO hiPSCs and PDE patient-derived hiPSCs, differentiated into astrocytes, to investigate the underlying mechanisms and potential adjunct treatments for PDE. Both *ALDH7A1* KO and patient astrocytes showed elevation of PDE biomarkers. Bulk RNA sequencing was used to explore underlying mechanisms, which revealed impaired oxygen response and increased oxidative stress levels. This was further validated by increased markers of oxidative stress including higher levels of LPO (4-HNE) as well as elevated OCRs in PDE astrocytes. Notably, targeting *AASS* in PDE astrocytes using CRISPR/Cas9 and AON-mediated downregulation significantly improved the metabolic phenotype and reduced OCRs, highlighting *AASS* reduction as a promising therapeutic approach for PDE.

Our phenotypic assessment of PDE astrocytes revealed differences between the *ALDH7A1* KO and PDE patient astrocytes. We found that *ALDH7A1* KO and PDE patient astrocytes showed a similar trend, but these phenotypes were milder in the patients compared to the *ALDH7A1* KO astrocytes, similar as has been described for another IMD^47^. We also observed significant differences between the three PDE patient astrocytes. For example, P2 astrocytes showed significantly higher PDE biomarker levels compared to P1 and P3 astrocytes, despite all patients sharing the same mutation, and P1 and P2 being siblings reducing the possible effects of variability due to genetic background. A key factor underlying the pronounced difference between the *ALDH7A1* KO and PDE patient astrocyte phenotypes is the complete loss of *ALDH7A1* expression in the *ALDH7A1* KO astrocytes due to the full knockout of the gene. In contrast, PDE patients harbor a missense variant that allows expression of the ALDH7A1 enzyme (Supplementary Figure 8), potentially retaining some residual enzymatic activity, although this remains uncharacterized for the PDE patient lines. In addition to its role in the lysine pathway, ALDH7A1 has been implicated in protecting against oxidative stress through the production of osmolytes as well as by the detoxification of aldehydes produced during LPO^27–29^. In line with our results, ALDH7A1 deficiency has previously been linked to ROS imbalance and subsequent oxidative stress^26^. This could also explain the more pronounced phenotype in the *ALDH7A1* KO astrocytes compared to the PDE patient astrocytes. The elevation of 8-Oxo-dG exclusively in *ALDH7A1* KO astrocytes, but not in any PDE patient-derived line, suggests that ALDH7A1 primarily protects against DNA oxidation, with a less pronounced role in mitigating LPO-related oxidative stress.

Increased levels of PA have been previously associated with elevated oxidative stress levels through increases in H_2_O_2_ for several other metabolic disorders^48,49^ and have been described to affect antioxidant capacity of the astrocytes^10^. In addition, accumulation of PA and P6C^38–40^ as well as increased oxidative stress^41–44^ are known to contribute to mitochondrial dysfunction. Our findings also show a correlation between elevated PDE biomarkers and dysregulated OCR (either decreased, indicating impaired mitochondrial function, or increased, suggesting potential compensatory upregulation), both of which are indicative of mitochondrial dysfunction. Interestingly, P1 and P3 showed increased 4-HNE but not P2, whereas P2 astrocytes showed highest levels of PA. However, ALDH7A1 levels were much higher in P2 compared to P1 and P3 (Supplementary Fig. 9). Increased ALDH7A1 levels, and therefore possibly increased antioxidant function might directly protect against the oxidative damage. The reason why ALDH7A1 levels are higher in P2 compared to the other patients remains unknown and requires further investigation. Altogether, the interplay between OCR, oxidative stress, and the accumulation of toxic lysine metabolites in PDE appears to be complex and bidirectional, where increased OCR can exacerbate oxidative stress, while oxidative stress and the buildup of metabolites like PA may impair mitochondrial function, leading to altered OCR. These dynamic interactions underscore the need for further investigation to better understand their combined role in disease progression and potential therapy.

One of the limitations of this study is the use of unrelated controls for comparison with PDE patient astrocytes. While these controls provided a necessary baseline, inherent genetic variability between unrelated individuals introduced increased variability in the assays, making it challenging to consistently achieve statistical significance across some parameters. To address this limitation, the inclusion of isogenic controls, such as PDE patient astrocytes with corrected mutations via CRISPR/Cas9, could serve as a more precise comparison. In general, metabolism is highly dynamic^50,51^, so the observed differences could also be dependent on the state of the cultured astrocytes. Although we did not observe visual differences between the patient-derived astrocytes it is possible that the astrocyte cultures have a different developmental trajectory affecting their metabolism.

Similar to a strategy adopted for GA1, another IMD of lysine metabolism, we targeted *AASS* as a way to rescue the PDE metabolic deficits. We developed two independent strategies to downregulate *AASS*; CRISPR/Cas9 genome editing to show proof-of-concept and a gapmer which potentially could be translated into a therapy for PDE patients. The main difference between these two strategies is that for the *ALDH7A1/AASS* DKO astrocytes, *AASS* is not present throughout development and therefore accumulation of direct byproducts caused by lysine catabolism are minimal. In contrast, in the patient astrocytes, the AASS enzyme and therefore metabolite accumulation, are present during development and *AASS* can only be inhibited post-development. Both approaches showed partial rescue of the PDE biomarkers and rescue of the OCR. In contrast to the P2 astrocytes, which did not exhibit increased oxidative stress at baseline, both DNA oxidation and LPO stress levels were rescued in the *ALDH7A1/AASS* DKO astrocytes. The rescue of oxidative stress in *ALDH7A1/AASS* DKO astrocytes, but not in G3-treated patient astrocytes, suggests that the observed oxidative stress in patient astrocytes might be unrelated to the lysine pathway. However, we cannot rule out that longer treatment durations or higher silencing effect might be necessary to achieve a measurable reduction of oxidative stress markers in PDE patient astrocytes. Still, the effects on the metabolic phenotype and OCR of the *ALDH7A1/AASS* DKO astrocytes and the G3-treated P2 astrocytes (around 50% remaining AASS protein levels) were relatively similar. This suggests that partial reduction in *AASS* levels (50%), instead of complete knock-down of *AASS*, is already sufficient for a partial rescue in PDE patient astrocytes, although there is variability across different PDE patients (Supplementary Fig. 7). Here we show proof-of-concept data for a potential therapeutic strategy to treat PDE. Further development of the molecule by optimizing the sequence, but also performing more *in vitro* and *in vivo*, as well toxicity and safety studies, are required to explore the full potential of this approach.

In conclusion, our study suggests that dysregulated OCR and increased oxidative stress contribute to the disease mechanism in PDE. Additionally, we provide the first evidence that substrate reduction through *AASS* inhibition could be a potential therapeutic approach for PDE. We demonstrated that partial reduction of AASS levels leads to a partial rescue of PDE biomarkers, elevated OCR and oxidative stress in PDE astrocytes, offering a promising avenue for future treatment strategies for PDE patients.

## Materials and methods

### hiPSC line information

In this study four independent control lines were used, of which C1, C2 and C3 are commercially available: C1 is derived via episomal reprogramming from fibroblasts of a 30-year-old Japanese male control (GM25256; from Coriell Institute, Camden, USA); C2 is derived from fibroblasts of a healthy 24-year-old female (UMGi020-A; from University Medical Center Goettingen; Goettingen; Germany) and C3 is derived from fibroblasts of a 40-year-old female control (HPSI0314i-hoik_1; from Cambridge BioResource; Cambridge; United Kingdom). C2 and C3 are both reprogrammed using Sendai viral vectors. C4 was generated via lentiviral reprogramming from fibroblasts of a 41-year-old female control.

Diagnosis of the PDE patients was based on both genetic and metabolic screening. All patients presented typical characteristics of PDE at the time of biopsy and all harbor the c.1279G>C (p.Glu427Gln) variant in ALDH7A1 in homozygosis. P1 and P2 are siblings, minimizing the effect of genetic background. The PDE patient hiPSC lines were created via episomal reprogramming of the patients-fibroblasts upon approval by the ethics committee and signed informed consent. Lines were fully validated as described elsewhere^23^. In addition, C1 was used to create a full knock-out of ALDH7A1 using CRISPR/Cas9 editing system. Generation and validation of this line has previously been described^24^. Accordingly, the ALDH7A1 KO line was used to create an additional knock-out of AASS using CRISPR/Cas9 resulting in a homozygous missense variant in intron 4 (c.388-640A>G). Two guide RNA were designed (TGATACAGCCTTCGAATCGGCGG/TCATAGAGGAGTACGGGTAGTGG) and cloned into pSpCas9(BB)-2A-Puro (PX459) V2.0 (Addgene, #62988) similarly as has been described for the ALDH7A1 KO line^24^.

### hiPSC-astrocyte differentiation

Astrocytes were differentiated according to a previously described protocol^25^. Briefly, hiPSCs were seeded upon dissociation with TrypLE (Gibco; 12604021) onto human recombinant laminin-521 (20 µg/mL; Biolamina; LN521-05 diluted in 1x dPBS++ (Gibco; 14040117)) pre-coated 6-well plates and cultured in Astrocyte medium (AM; ScienCell®; 1801) supplemented with RevitaCell™ (Gibco; A2644501) and Primocin at 37 °C/5% CO2. The next day AM supplemented with Primocin was refreshed to remove dead cells and to withdraw RevitaCell™ from the medium. AM supplemented with Primocin was refreshed every other day and at 100% confluency the astrocytes were split by dissociation using TrypLE. Accordingly, the cell pellet was resuspended in AM supplemented with RevitaCell™ and Primocin and all cells were transferred into either a T25 flask (Corning; 430372) during the first passage or a T75 flask (Corning; 430641U) during the second passage, followed by full change of AM supplemented with Primocin the day after. Hereafter, the astrocytes were split at 90-100% confluency at 1:3 onto T75 flasks and cultured in AM with Primocin but in the absence of RevitaCell™. After five weeks of differentiation, medium changes were performed only twice a week. The cells were imaged using the Invitrogen™ EVOS™ XL Core Configured Cell Imager.

### hiPSC maintenance

hiPSCs were cultured on Geltrex-coated (Gibco; A1413301) 6-well plates (Corning; 353046) in Essential 8TM Flex Medium kit (Gibco; A2858501) supplemented with Primocin (0.1 μg/ml; Invivogen; ANT-PM-2) at 37 °C/5% CO2. Medium was refreshed every two to three days, and at around 80% confluency the hiPSCs were passaged using ReLeSR (Stemcell Technologies; 100-0483) upon washing in Dulbecco’s phosphate-buffered saline (dPBS; Gibco; 14190169).

### Immunocytochemistry

During fixation, the cells were incubated with 4% paraformaldehyde (Sigma Aldrich; 441244)/4% sucrose (Sigma; S7903) for 15 min and accordingly washed three times with PBS (1x; Sigma; P5493). The cells were blocked for 1 h with blocking buffer (BB; 5% normal donkey serum (Jackson Immuno Research; 017-000-121), 5% normal goat serum (Invitrogen; 10189722), 5% normal horse serum (Gibco; 26050070), 0.1% D-lysine (Sigma-Aldrich; L8021), 1% glycine (Sigma-Aldrich; G7126), 1% BSA (Sigma-Aldrich; A-6003) and 0.4% Triton X-100 (Sigma-Aldrich; T8787) in PBS) at room temperature (RT). Primary and secondary antibodies were diluted in BB and incubated at 4 °C overnight or at RT for 1 h, respectively. Cells were incubated for 10 min at RT with Hoechst (Thermo Scientific; 62249) diluted in PBS to stain for the nuclei of the cells. Accordingly cells were mounted using DAKO fluorescent mounting medium (DAKO; S3023). The following antibodies were used: Rabbit anti-Vimentin (1:300; Abcam; ab92547), Rabbit anti-GFAP (1:300; Sigma-Aldrich; AB5804), Mouse anti-CD44 (1:300; Invitrogen; MA5-13890), Mouse anti-ALDH1L1 (1:300; Novus Biologicals; NBP2-50045), Rabbit anti-GLUD1 (1:300; Invitrogen; PA5-28301), Mouse anti-8-Oxo-dG (1:500; R&D systems; 4354-MC-050), Mouse anti-4-HNE (1:500; R&D systems; MAB3249), Goat anti-Rabbit Alexa Fluor 568 (1:1000; Invitrogen; A11011), Goat anti-Mouse Alexa Fluor 488 (1:1000; Invitrogen; A11029). Zeiss Axio Imager Z1 was used to image the samples.

### NGMS sample preparation and analysis

To measure the PDE biomarkers within the astrocytes, cells were prepared by dissociation using TrypLE, plated onto 12-well plates (Corning; 353043) and cultured in AM at 37 °C/5% CO2. One day prior harvesting, AM was fully refreshed. Biological triplicates of each sample were prepared for NGMS analysis according to a previously described protocol^52^. Briefly, the astrocytes were washed twice with RT sample buffer, consisting of ammonium carbonate (75 mM; Honeywell; 207861-500G) diluted in Ultrapure water and adjusted pH of 7.2– 7.4 using acetic acid. Immediately upon washing with sample buffer, the cells were frozen in liquid nitrogen on the culture plates. Metabolite extraction from the frozen cells was done by 2 min incubation with 250 μL cold extraction buffer (2:2:1 (v/v/v) methanol (Honeywell;34860–2.5L) : acetonitrile (Biosolve; 0001207802BS) : Ultrapure water) followed by another 3 min incubation with new extraction buffer. The two extracts were pooled and centrifuged at 14,000 x g for 3 min at 4 °C. Accordingly, the supernatant was dried using a vacuum centrifuge (SpeedVac, Thermo Fisher Scientific) and the samples were stored at −80 °C until further use. Metabolites were reconstituted in 80 μL Ultrapure water/0.1% HCOOH and analyzed using an ultra-high-performance liquid chromatography-quadrupole time of flight mass spectrometry (UHPLC-QTOF-MS54). Analyses were performed using an Agilent 1290 UHPLC system coupled to an Agilent 6545 QTOF mass spectrometer, equipped with a dual electrospray ionization (ESI) source. Each sample was run in duplicate in both negative and positive ionization modes. Two microliters of extracted metabolite sample was injected onto an Acquity HSS T3 (C18, 2.1 × 100 mm, 1.8 μm) column (Waters) operating at 40 °C. Each analytical batch included astrocyte samples, analytical quality control (QC) samples and a pooled sample of all astrocyte samples to check for integrity of the automated data analysis pipeline. The biological triplicates of all samples were measured in duplicate. To correct for possible run-order influence on signal intensities, the technical duplicates were analyzed in antiparallel run order. In addition, eight random astrocytes samples were injected at the start of each analytical batch to condition the analytical platform. For analysis, relative peak area was extracted and accordingly normalized for the peak area of phenylalanine.

### RNA sequencing

#### RNA-seq library preparation

Astrocyte cultures of C1, C4, ALDH7A1 KO, P1 and P3 were prepared for RNA sequencing. Biological triplicates of all astrocyte cultures were seeded onto 6-well plates in AM and cultured at 37 °C/5% CO2. At confluency, the astrocytes were washed with ice-cold PBS twice and were then collected using DNA/RNA shieldTM (Zymo Research; #R1200-125). Subsequently, RNA extraction was performed utilizing the Quick-RNA Microprep kit (Zymo Research, #R1051) following the manufacturer’s protocol. The quality of the extracted RNA was evaluated using Agilent’s Tapestation system, with RNA Integrity Number (RIN) values falling within the range of 7.3 to 9.6. Next, cDNA libraries were prepared using the NEBNext Ultra II Directional RNA Library Prep Kit, followed by sequencing of paired-end reads on an Illumina NovaSeq 6000 platform at GenomeScan B.V. Leiden.

#### RNA-seq data processing

We used Fastp^53^ to eliminate PolyG artifacts and clip adapters, including a list of adapter sequences currently used by Illumina. Subsequently, we mapped the reads to the GRCh38 human reference genome using HISAT2. As the library was reversely stranded, we set-rna-strandness = RF. The resulting SAM files were sorted and indexed using SAMtools^54^. Next, we performed UMI deduplication with UMI-tools^55^, and feature counting was done with Subread’s featureCounts^56^.

#### RNA-seq analysis

Raw count matrices were loaded in R version 4.2.1. When Ensemble IDs mapped to the same gene symbols, we considered only the IDs with the highest expression per sample. Counts belonging to the Y chromosome were excluded. We filtered out lowly expressed genes by keeping genes with expression values higher than 0.5 transcripts per million reads in at least three samples. The remaining counts were then transformed into a DGEList using the DGEList function from edgeR package^57^. Subsequently, voom normalization of limma package61 was applied, including normalization factors determined by edgeR’s calcNormFactors. Principal component analysis (PCA) was conducted on voom normalized and scaled data using prcomp function. Differential expression analysis was carried out for ALDH7A1 KO and Ctrl astrocytes. Additionally, comparisons were made between the PDE patients and the two control cell lines. For both cases, we normalized using voom and factoring based on condition. Subsequently, limma’s functions lmFit, makeContrasts, contrasts.fit, and eBayes were used to estimate the condition coefficient and its corresponding p-value for each gene. Genes were defined as differentially expressed if they exhibited a Benjamini-Hochberg (BH)-adjusted p-value below 0.05, coupled with a Log2 fold change exceeding 0.58.

Enrichment analysis was separately conducted for differentially expressed genes (DEGs) in ALDH7A1 KO vs control and PDE patients vs controls comparisons, using the go function from the R package gprofiler2 v0.2.1. To manage redundancy among enriched gene ontology terms (including biological processes, cellular components, or molecular functions), we performed clustering analysis and aggregated terms with high semantic similarity with the functions calculateSimMatrix and reduceSimMatrix from the rrvgo v1.2.0 R package, setting threshold=0.7.

#### Seahorse

OCR was assessed using the Agilent Seahorse XF Cell Mito Stress Test Kit (Seahorse Bioscience). Astrocytes were seeded at a density of 8,000 cells per well for baseline experiments and 4,000 cells per well for gapmer experiments, in AM supplemented with Primocin and RevitaCellTM at 37 °C/5% CO2. The day after plating, AM supplemented with Primocin was fully refreshed to withdraw RevitaCellTM from the medium. One hour before the assay, AM was replaced with Agilent Seahorse XF Base Medium supplemented with 10 mM glucose, 1 mM sodium pyruvate and 200 mM L-glutamine, and the cells were incubated at 37 °C without CO2. During the recording, basal oxygen consumption was measured four times, followed by three measurements after each addition of 1 μM oligomycin A, 1 to 4 μM carbonyl cyanide 4-trifluoromethoxy phenylhydrazone (FCCP), 0.5 μM rotenone and 0.5 μM antimycin A. Each measurement cycle consisted of 3 min of mixing, 3 min of waiting, and 3 min of measuring. The OCR values were normalized for protein concentration to account for differences in cell number and cell size. To determine the protein concentration for each well, the Pierce™ BCA protein assay was used according to the manufacturer’s instructions (Thermo Scientific 23225). The most optimal FCCP concentration (e.g. maximal respiration) was determined for each line and used for further analysis.

#### Oxidative stress assay

To measure oxidative stress in astrocytes, immunostainings for both 8-Hydroxy-2′-deoxyguanosine (8-Oxo-dG) and 4-Hydroxynonenal (4-HNE) were performed. 8-Oxo-dG is an oxidized derivative of deoxyguanosine, one of the major products of DNA oxidation and therefore considered a measure for oxidative stress^58^. 4-HNE is an α,β-unsaturated hydroxyalkenal that is produced during LPO in cells. Increased 4-HNE is therefore also associated with increased oxidative stress^59^. For both assays the astrocytes were plated either onto glass coverslips (Epredia; 631-0713) or on 96-well culture plates (Greiner; 655090). For baseline characterization the cells were fixed at 70-95% confluency, around 3-4 days after seeding. The AON-treated astrocytes were fixated seven days after AON-delivery. Immunostainings for 8-Oxo-dG and 4-HNE were performed as described previously.

We imaged at a 20x magnification using the Zeiss Axio Imager Z1. To compare signal intensities between different experimental conditions, all conditions within a batch were acquired with the same settings. Fluorescent signals were quantified using FIJI software. The mean intensity per image was calculated and corrected for the total area of the cells in the respective image. The total area was based on the astrocyte specific cytoskeleton marker vimentin. Fold Change (FC) intensity was measured for each well relative to the averaged intensity of the merged controls for baseline characterization or relative to the NT condition for the gapmer experiments.

#### AON design

The AONs used in this study were designed following the guidelines for splice switching antisense oligonucleotides in terms of region accessibility^60^. To analyze the accessibility of the target RNA secondary structure, Mfold software version 6.4 was used. The sequence properties, including length, GC content, and Tm of the AONs, were calculated using OligoCalc online software version 3.27. Customized oligonucleotides were purchased from Eurogentec (Liege, Belgium). The gapmer and sense oligonucleotide (SonG) control contained a core DNA sequence of 12 nucleotides with a phosphorothioathe (PS) backbone, flanked by 4 RNA-nucleotide LNA/PS wings (G3 sequence: UCCU-GAAGAATGCCAC-CAGC; SonG sequence: CGAC-CACCGUAAGAAG-UCCU). The lyophilized gapmer and SonG were resuspended in sterile PBS to a working concentration of 100 µM. For the other AONs (ssAONs and gapmers) designed in this study see Supplementary Table 1.

#### Gapmer delivery

Astrocytes were seeded onto 12-well culture plates in AM. The day after, the gapmer was delivered at two different concentrations (0.05 and 0.5 µM), whereas the SonG was delivered only at the highest concentration (0.5 µM). One well was kept untreated (NT) in parallel. The gapmer was delivered using FuGENE HD reagent (Promega, Madison, WI, USA) as previously described64. AM was refreshed every two to three days (without the gapmer). Seven days after gapmer delivery the astrocytes were either used for seahorse assay, fixated for oxidative stress assays or collected for RNA analysis or western blot.

#### RT-qPCR

Total RNA was extracted from the cells using the NucleoSpin RNA Mini Kit (Macherey-Nagel; MN 740955.250) following the manufacturer’s instructions, with the exceptions of omitting β-mercaptoethanol and extending the incubation time with rDNase to 30 min instead of 15 min. Subsequently, the RNA was reverse-transcribed into cDNA utilizing the iScript cDNA Synthesis Kit (Bio-Rad; 1708891). For quantitative PCR (qPCR), GoTaq qPCR Master Mix (Promega; A6002) was used. The qPCR was conducted on the 7500 Fast Real-Time PCR System (Applied Biosystems) with the following cycling conditions: denaturation at 95°C for 2 min, followed by 40 cycles of 30 sec at 95°C and 30 sec at 60°C, and a melting curve stage of 15 sec at 95°C, 30 sec at 60°C, and 15 seconds at 95°C. All samples were prepared in triplicate, and outliers were identified if a value deviated by more than 0.5 Ct from the other two values within the technical triplicate. The relative mRNA expression was determined using the 2−ΔΔCt method, normalized to the housekeeping gene GUSB. Primers are listed in Supplementary Table 1.

#### Western blot

To lyse the cells, the medium was aspirated, and the cells were washed with PBS. Subsequently, lysis buffer (composed of RIPA buffer (pH 7.5; 50 mM Tris-HCl (Invitrogen; 15567027), 150 mM NaCl (Sigma-Aldrich; S5886), 1 mM EDTA (Sigma-Aldrich; 03690), and 1% Triton X-100 in PBS), supplemented with protease inhibitors (cOmplete Mini; Roche; 11,836,153,001) was added to the cells. Prior to blotting, the protein concentration was determined using PierceTM BCA protein assay according to the manufacturer’s instructions (Thermo Scientific; 23227). Subsequently the samples were loaded with an equal amount of protein (ranging from 5 to 15 μg) onto 4-15% Mini-PROTEAN® TGX Stain-Free™ Protein Gels (Bio-Rad; 4568084) for protein separation through SDS-PAGE (Bio-Rad; 30 min at 200 V). Accordingly, the separated proteins were transferred to nitrocellulose membranes (Bio-Rad, #1704158) using the Trans-Blot Turbo Transfer System (Bio-Rad; 7 min at 25 V and constant 1.3 A). Membranes were blocked in 5% non-fat milk (Santa-Cruz; sc-2325) in TBS (Millipore; 524750-1EA)/0.1% tween (Merck; 8.22184.0500) (TBS-T) for 1 h at RT and incubated overnight at 4 ◦C with the respective primary antibodies diluted in 5% non-fat milk in TBS-T. The following primary antibodies were used: rabbit anti-AASS (1:1000; Prestige antibodies; HPA020734-100UL), rabbit anti-ALDH7A1 (1:500; Invitrogen; PA5-54750), rabbit anti-GAPDH (1:2000; Cell Signaling Technology, 2118). Horseradish peroxidase-conjugated goat anti-rabbit antibodies (1:50,000; Invitrogen, G21234) were used for visualization.

#### Statistical analysis

Statistical analysis was performed using Graphpad Prism (version 10 for Windows, GraphPad Software). We first determined whether data were normally distributed using the D’Agostino & Pearson test, Anderson-Darling test, Shapiro-Wilk test and Kolmogorov-Smirnov test. When only two conditions were compared, unpaired t test was used. When data of only two conditions were not normally distributed we performed Mann-Whitney test. To test statistical significance for three or more conditions (either different cell lines or treatments), one-way ANOVA combined with Sidak’s multiple comparison tests was used. If these data were not normally distributed, we performed Kruskal-Wallis test combined with Dunn’s testing for multiple comparisons. Data are shown as mean and standard deviation (SD) in bar diagrams unless stated otherwise. Results with P values lower than 0.05 were considered as significantly different (*), P < 0.01 (**), P < 0.001 (***), P < 0.0001 (****). Details about statistics are reported in Supplementary Table 1.

## Data availability

The datasets generated during the current study are available from the corresponding author on reasonable request. Source data underlying Figs. 1–5 and Supplementary Figs. 1–9 is available in Supplementary Table 1. The GEO accession number for the RNA-seq data in this paper is GSE287216. Access Token: clsxaiwcvnanrch

## Acknowledgements

We gratefully acknowledge the entire CHARLIE consortium for the constant feedback and support. In particular we would like to acknowledge Prof. Ron Wevers for insightful discussions about the metabolomic measurements. We would also like to thank Dr. Werner Koopman and Dr. Merel Adjobo-Hermans for their assistance with identifying proper oxidative stress readouts. Furthermore, we want to acknowledge all members of the Nadif Kasri lab, and the Collin & Garanto lab for the continuous help and support.

This study has been initiated by the CHARLIE consortium (EJPRD grant no: 825575 awarded to Clara D.M. van Karnebeek and Nael Nadif Kasri). Imke M.E. Schuurmans was supported by an internal Radboudumc PhD grant provided by the Radboud Institute for Molecular Life Sciences (awarded to Clara D.M. van Karnebeek and Alejandro Garanto) and the Catalyst Grant form United for Metabolic Diseases (UMD-CG-2023-001) which is financially supported by MetaKids (awarded to Alejandro Garanto, Imke M.E. Schuurmans, Clara D.M. van Karnebeek and Nael Nadif Kasri).

## Author contributations

C.v.K, N.N.K and A.G. conceived and supervised the study. I.S., C.v.K., B.L., N.N.K and A.G. (partially) designed the experiments. I.S., U.E., S.P., R.M., G-J.S., S.v.K. and A.O. performed or assisted during the experiments. C.v.K., N.N.K. and A.G. provided resources. I.S., U.E., M.A., S.P., R.M., G-J.S., S.v.K., A.O., H. A-S., D.L., C.v.K., N.N.K. and A.G. performed or assisted during data analysis and intepretation. I.S., N.N.K. and A.G. wrote the paper. U.E., M.A., S.P., R.M., G-J.S., S.v.K., A.O., D.L., B.L. and C.v.K. edited the paper.

