## Supplementary Figures for "Targeting AASS improves neurotoxicity and mitochondrial function in astrocyte models for pyridoxine dependent epilepsy"

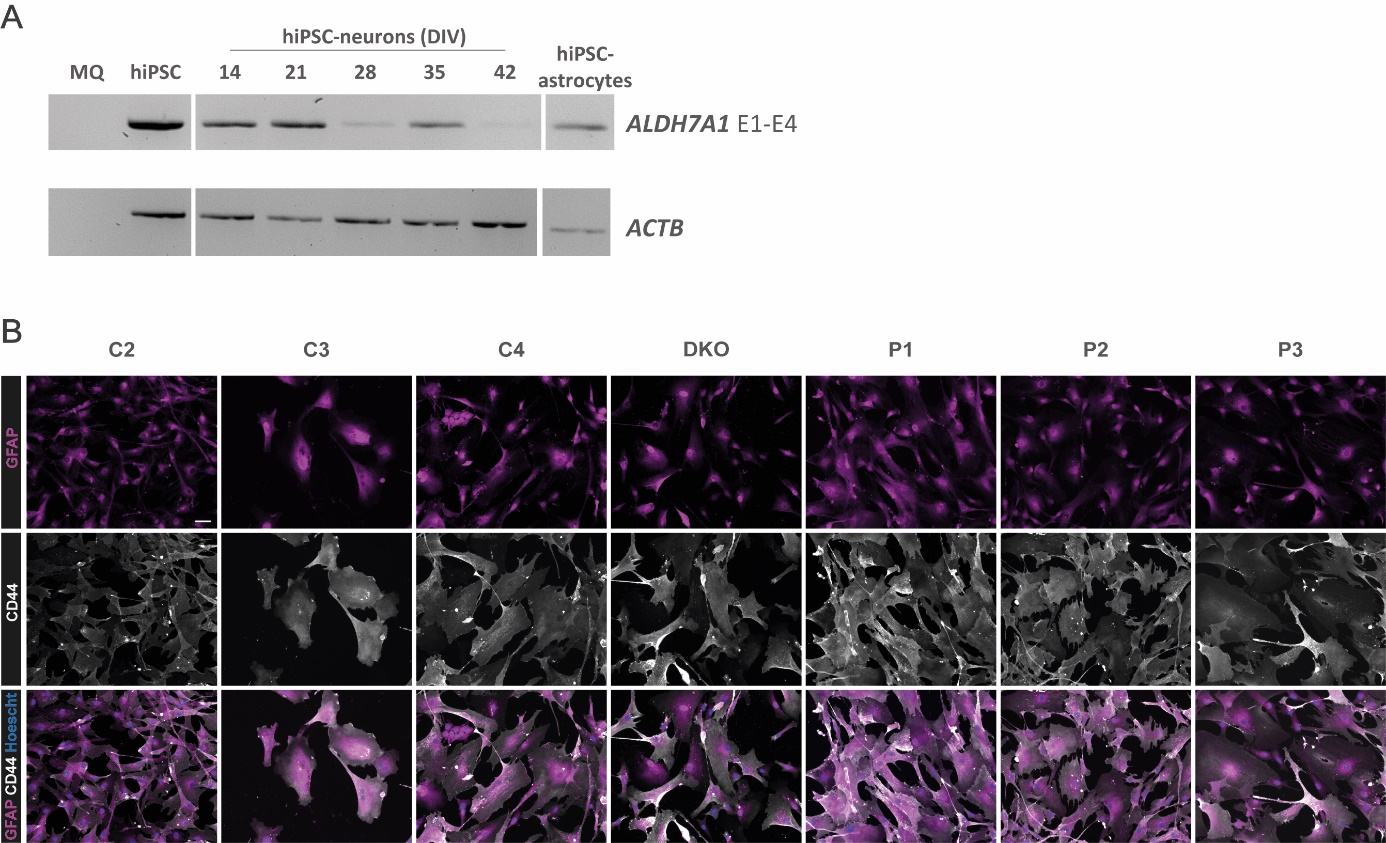

**Supplementary Figure 1. Characterization of all PDE astrocytes. A.** Expression of *ALDH7A1* relative to *ACTB* by regular PCR in hiPSCs, hiPSC-derived neurons from 14, 21, 28, 35 and 42 days in vitro (DIV) and hiPSC-derived astrocytes. **B.** Representative images of immunostaining of GFAP (magenta), CD44 (white), GLUD1 (magenta) and ALDH1L1 (white) in DIV 35 astrocytes from C2, C3, C4, *ALDH7A1/AASS* KO, P1, P2 and P3. All pictures were taken at the same magnification (scale bar = 50 µm).

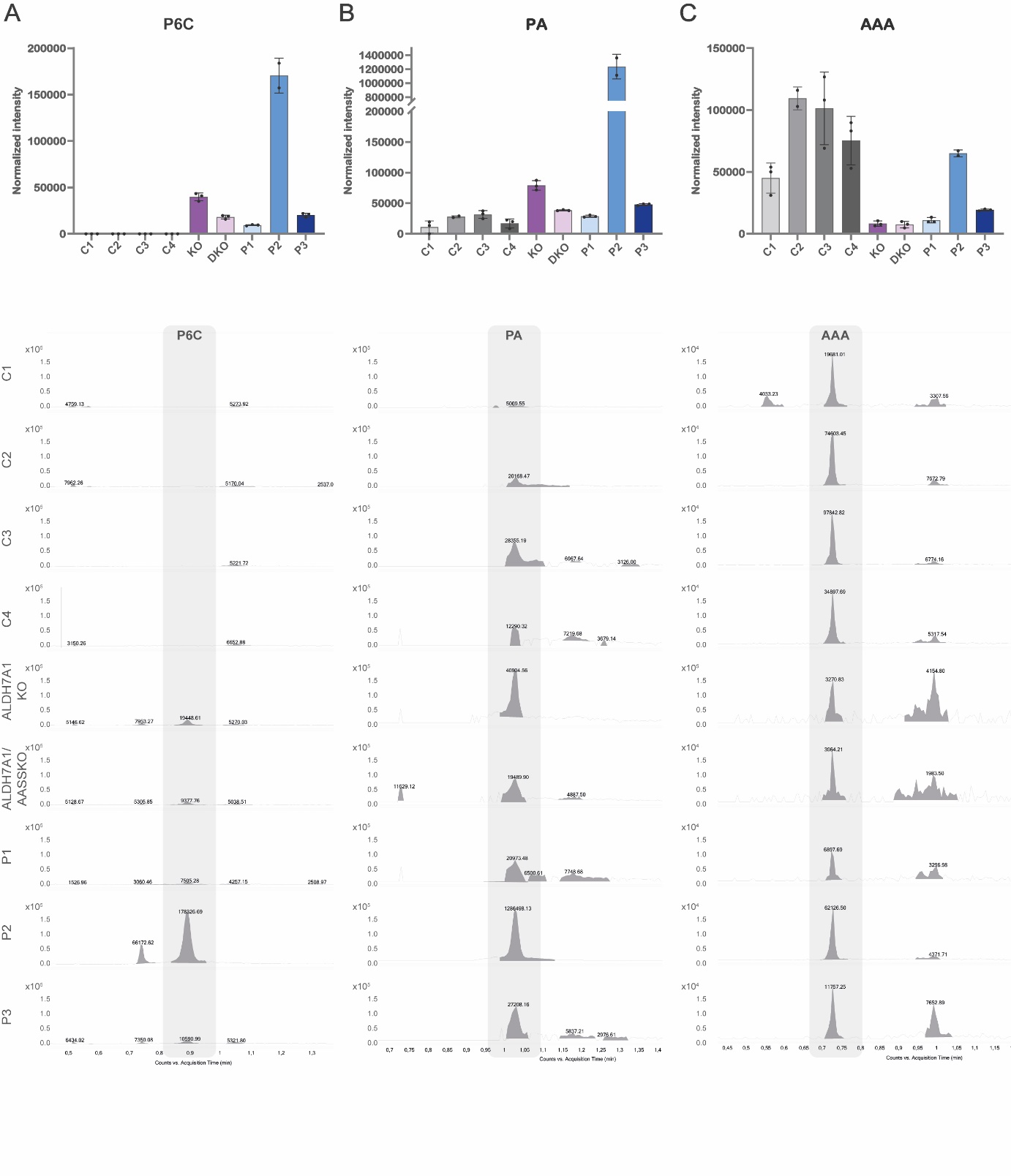

**Supplementary Figure 2. Metabolic characterization of all PDE astrocytes. A.** The graph shows the normalized intensity of P6C relative to phenylalanine measured via NGMS in astrocytes from control, *ALDH7A1* KO (KO), *ALDH7A1/AASS* DKO (DKO) and PDE patients. *n* = 3 for C1; *n* = 3 for C2; *n* = 3 for C3; *n* = 3 for C4; *n* = 3 for KO; *n* = 3 for DKO; *n* = 3 for P1; *n* = 2 for P2; *n* = 3 for P3. For one sample of the biological triplicate of each line the P6C peak at the correct retention time (indicated by grey box) is shown for all lines, including the relative peak intensity. **B.** The graph shows the normalized intensity of PA relative to the housekeeping metabolite phenylalanine measured via NGMS in astrocytes from control, KO, DKO and PDE patients. *n* = 3 for C1; *n* = 3 for C2; *n* = 3 for C3; *n* = 3 for C4; *n* = 3 for KO; N = 3 for DKO; *n* = 3 for P1; *n* = 2 for P2; *n* = 3 for P3. For one sample of the biological triplicate of each line the PA peak at the correct retention time (indicated by grey box) is shown for all lines, including the relative peak intensity. **C.** The graph shows the normalized intensity of AAA relative to the housekeeping metabolite phenylalanine measured via NGMS in astrocytes from control, KO, DKO and PDE patients. *n* = 3 for C1; *n* = 3 for C2; *n* = 3 for C3; *n* = 3 for C4; *n* = 3 for KO; *n* = 3 for DKO; *n* = 3 for P1; *n* = 2 for P2; *n* = 3 for P3. For one sample of the biological triplicate of each line the AAA peak at the correct retention time (indicated by grey box) is shown for all lines, including the relative peak intensity.

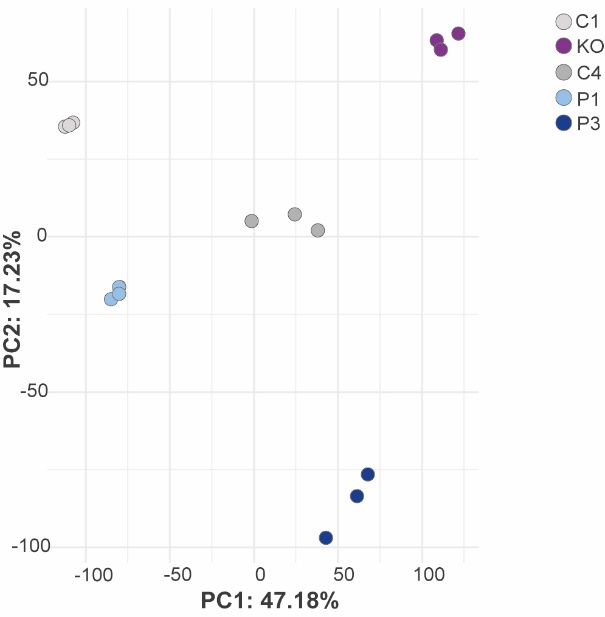

**Supplementary Figure 3. Principal component plot of RNA sequencing.** Principal component (PC) plot of RNA sequencing data representing biological triplicates of DIV 35 astrocytes derived from *ALDH7A1* KO (KO), P1, P3 and C1 and C4.

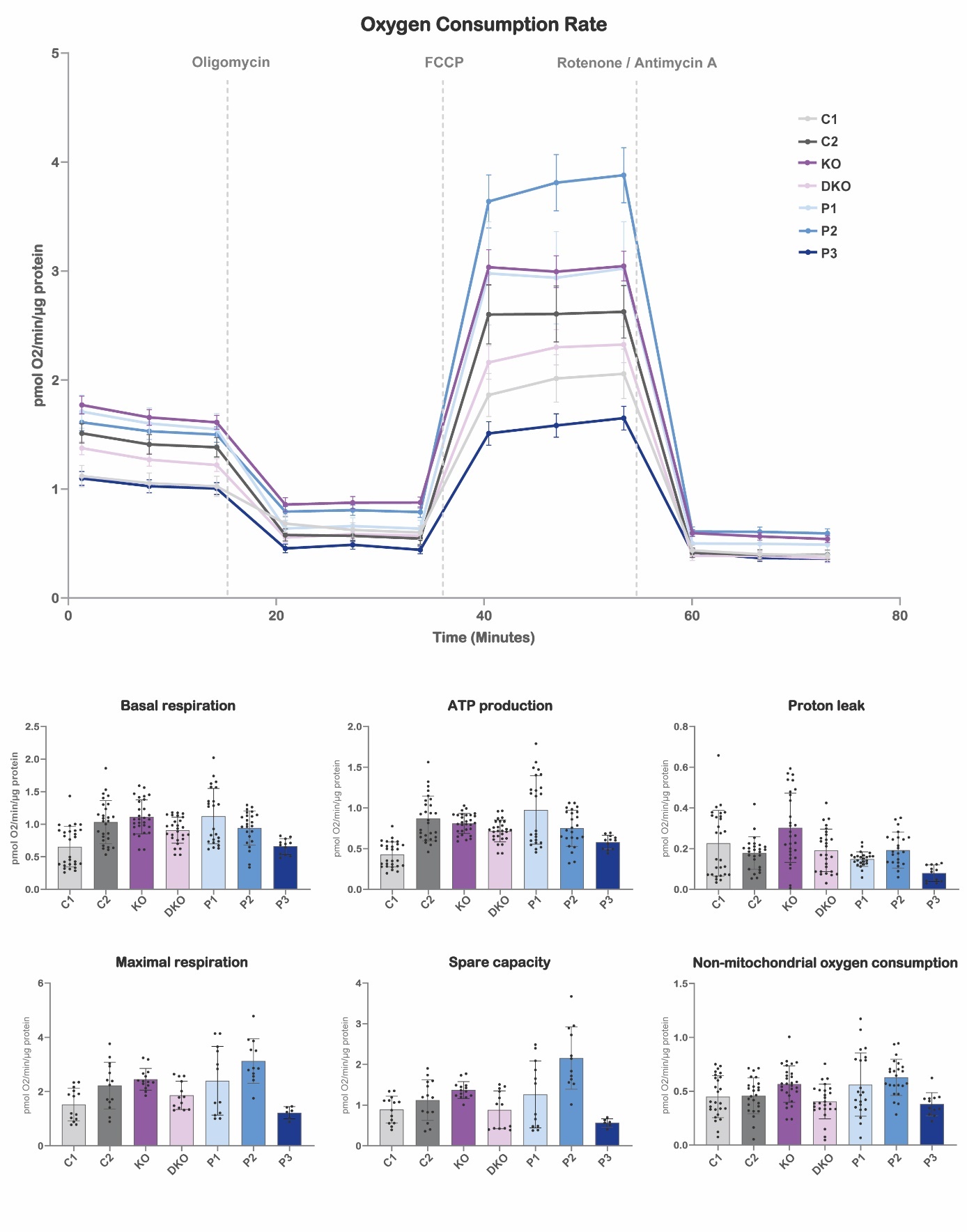
 **Supplementary Figure 4. Seahorse assay and immunostainings for oxidative stress metabolites.** OCR plot as well as Basal respiration (BR), ATP production (AP), proton leak (PL), maximal respiration (MR), spare capacity (SC) and non-mitochondrial oxygen consumption (NMOC) represented in bar graphs are shown for DIV 35 astrocytes from C1, C2, *ALDH7A1* KO (KO), *ALDH7A1/AASS* DKO (DKO) and PDE patients lines. For BR, AP, PL and NMOC: *n* = 28/2 for C1; *n* = 27/2 for C2; *n* = 29/2 for KO; *n* = 27/2 for DKO; *n* = 24/2 for P1; *n* = 24/2 for P2; *n* = 12/1 for P3. For MR and CP: *n* = 14/2 for C1; *n* = 14/2 for C2; *n* = 14/2 for KO; *n* = 13/2 for DKO; *n* = 12/2 for P1; *n* = 12/2 for P2; *n* = 6/1 for P3.

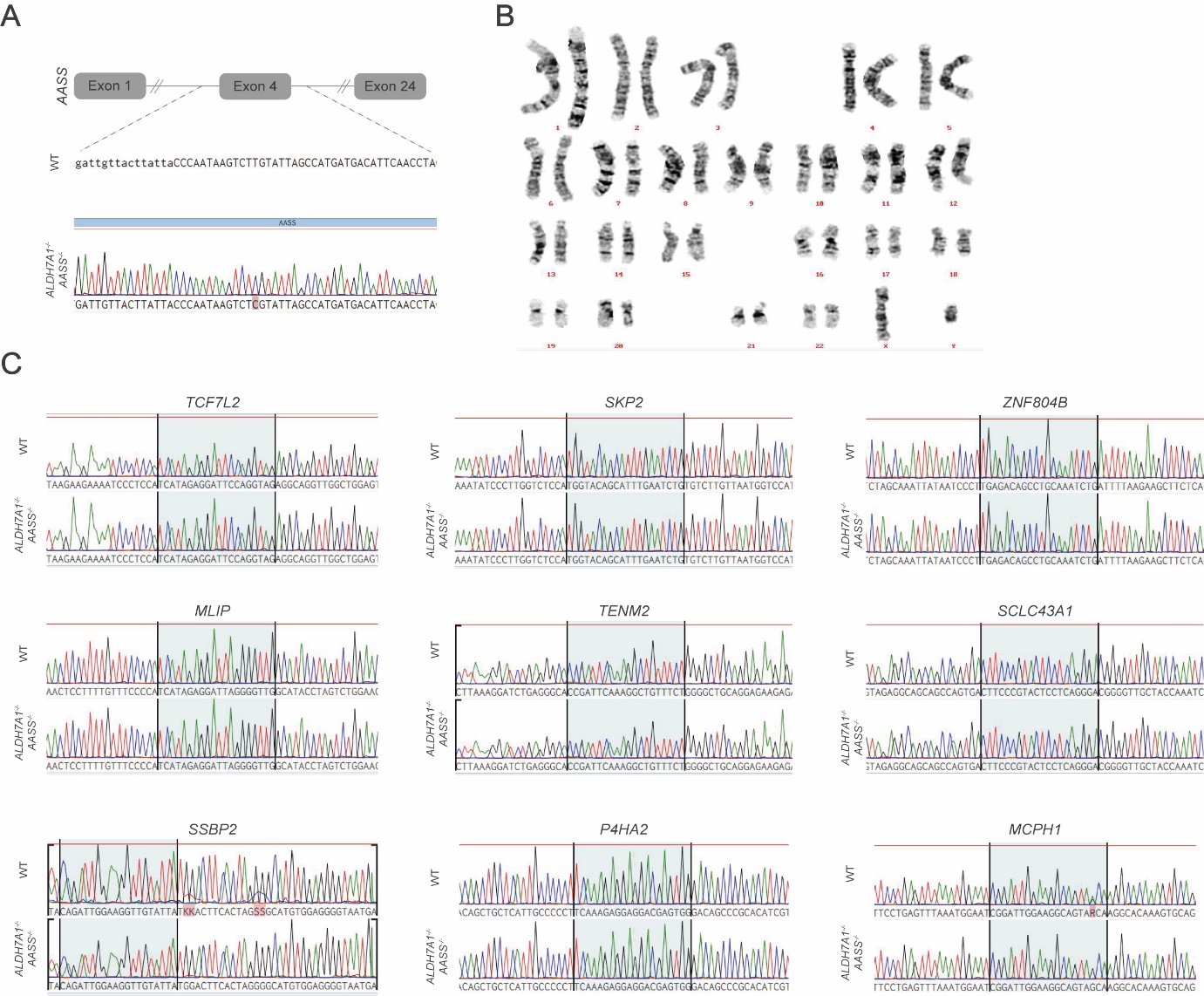

**Supplementary Figure 5. Generation of *ALDH7A1/AASS* DKO hiPSC line.** **A.** Schematic overview of CRISPR/Cas9 editing to create *ALDH7A1/AASS* DKO hiPSC line including chromatograms of sequencing results. **B.** Normal karyotype of *ALDH7A1/AASS* DKO hiPSC. **C.** All predicted off-target sites have been sequenced and no mutations were detected.

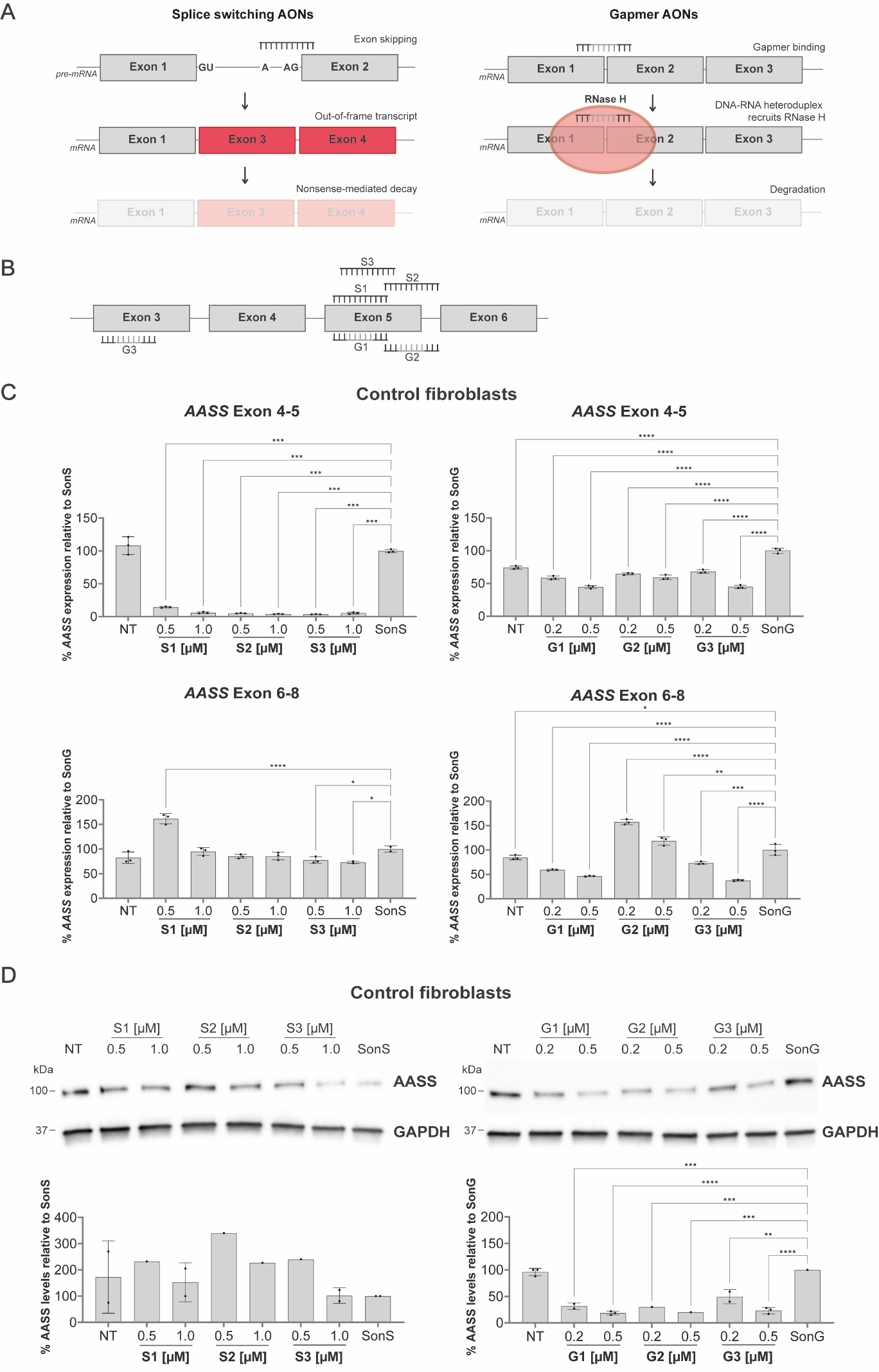
**Supplementary Figure 6. Screening of AONs targeting *AASS* in fibroblasts. A.** Schematic representation of two AON-strategies to target *AASS*; splice-switching AONs (ssAONs) and gapmers. **B.** Overview of the three designed ssAONs (top) and three gapmers (bottom). **C.** Relative expression of the regions exon 4-5 and exon 6-8 of *AASS* normalized for *GUSB* by RT-qPCR in control fibroblasts four days upon AON-delivery. ssAONs were transfected at 0.5 and 1.0 µM, gapmers were transfected at 0.2 and 0.5 µM, SonS was transfected at 1.0 µM and SonG was transfected at 0.5 µM**.** Data represents the percentage of remaining *AASS* expression relative to the SonG condition. **D.** Semi-quantification of AASS protein levels relative to GAPDH and representative (cropped) western blot of control fibroblasts four days upon AON-delivery. ssAONs were transfected at 0.5 and 1.0 µM, gapmers were transfected at 0.2 and 0.5 µM, SonS was transfected at 1.0 µM and SonG was transfected at 0.5 µM. Data represents the percentage of remaining AASS expression relative to the SonG condition. For exact *n* per experiment, per condition see Supplementary Table 1.

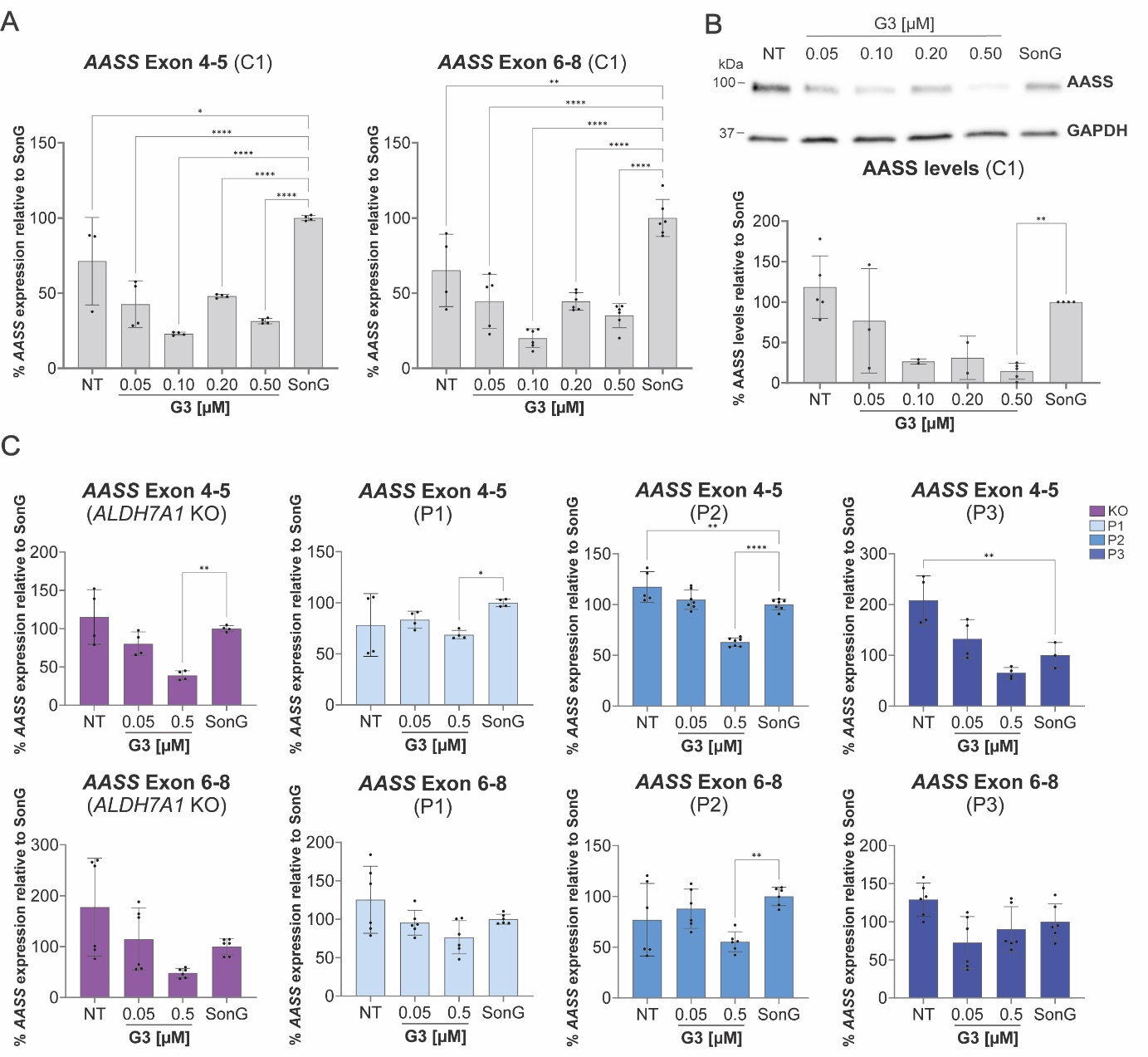
**Supplementary Figure 7. Screening of gapmers targeting *AASS* in control and PDE patient-derived astrocytes. A.** Relative expression of the regions exon 4-5 and exon 6-8 of *AASS* by qPCR in C1 astrocytes seven days upon G3-delivery at concentrations ranging from 0.05 - 0.5 µM and a sense oligonucleotide (SonG) control at 0.5 µM. Expression of genes was normalized against *GUSB*. Data represents the percentage of remaining *AASS* expression relative to the SonG condition. **B.** Semi-quantification of AASS protein levels relative to GAPDH and representative (cropped) western blot of C1 astrocytes seven days upon delivery G3 at concentrations ranging from 0.05 - 0.5 µM and SonG at 0.5 µM. Data represents the percentage of remaining AASS levels relative to the SonG condition. **C.** Relative expression of the regions exon 4-5 and exon 6-8 of *AASS* by qPCR in *ALDH7A1* KO, P1, P2 and P3 astrocytes seven days upon G3-delivery at 0.05 / 0.5 µM concentrations and SonG at 0.5 µM. Expression of genes was normalized against *GUSB*. Data represents the percentage of remaining *AASS* expression relative to the SonG condition. For exact *n* per experiment, per condition see Supplementary Table 1.

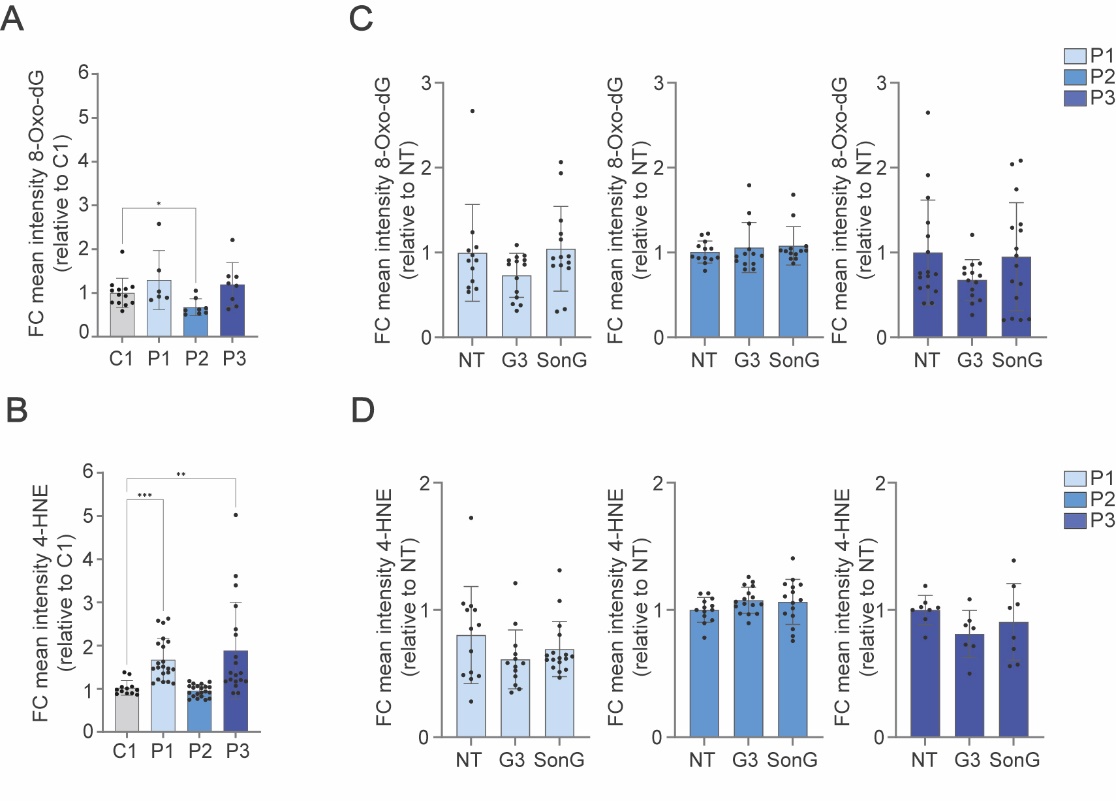
**Supplementary Figure 8. Oxidative stress upon gapmer treatment in PDE patient-derived astrocytes. A.** FC of mean intensity of 8-Oxo-dG per well relative to average intensity of C1 shown for P1 NT, P2 NT and P3 NT. **B.** FC of mean intensity of 4-HNE per well relative to average intensity of C1 shown for P1 NT, P2 NT and P3 NT. **C.** FC of mean intensity of 8-Oxo-dG per well relative to average intensity of corresponding NT shown for NT, G3 and SonG conditions in P1, P2 and P3 astrocytes. **D.** FC of mean intensity of 4-HNE per well relative to average intensity of corresponding NT shown for NT, G3 and SonG conditions in P1, P2 and P3 astrocytes. For exact *n* per experiment, per condition see Supplementary Table 1.

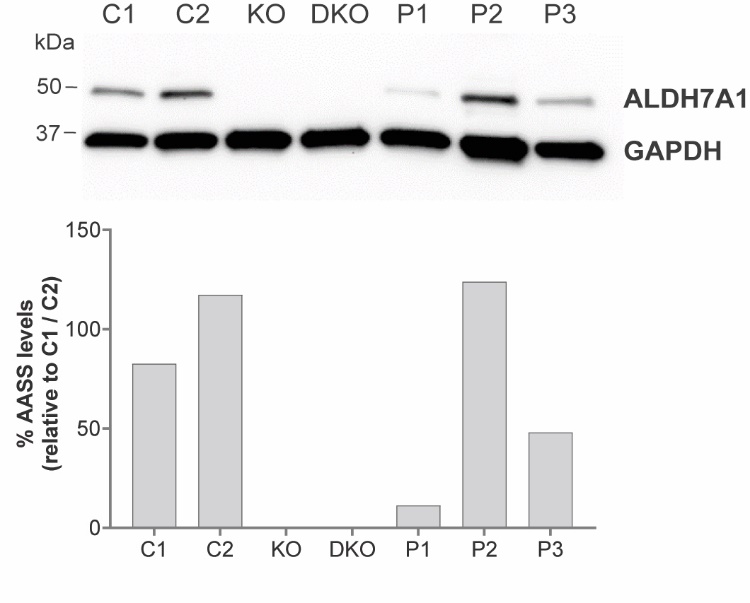

**Supplementary Figure 9. ALDH7A1 expression in the different astrocyte lines.** Semi-quantification of ALDH7A1 protein levels relative to GAPDH and (cropped) western blot of C1, C2, *ALDH7A1* KO (KO), *ALDH7A1/AASS* DKO (DKO), P1, P2 and P3 astrocytes. Data represents the percentage of remaining ALDH7A1 expression relative to the average of C1 and C2.

**Supplementary Table 1.** List of primers, clinical data on PDE patients and statistical information.
